# Rise-and-fall dynamics reveal a molecular and cellular vulnerability axis in prion-like α-synuclein propagation

**DOI:** 10.64898/2026.03.27.714785

**Authors:** Christoffer G. Alexandersen, Julia K. Brynildsen, Alice Prigent, Massimiliano Tamborrino, Anastasia Mantziou, Kevin Kurgat, Michael X. Henderson, Dani S. Bassett

## Abstract

The spread of misfolded proteins through neuronal circuits is a defining feature of neurodegenerative disease, yet the dynamics underlying this process remain poorly understood. Most studies rely on sparsely sampled datasets that capture spatial patterns of pathology but not their temporal evolution. Here, we analyze longitudinal histopathology measurements of *α*-synuclein pathology across hundreds of brain regions in a mouse model of Parkinson’s disease. Network-based dynamical modeling shows that regional pathology does not simply accumulate but instead follows rise-and-fall trajectories across the brain. The inferred parameter landscape reveals a one-dimensional vulnerability axis along which regions with stronger rise-and-fall dynamics have elevated energy metabolism and greater monoaminergic neuronal composition. This vulnerability axis replicates in an independent histopathological dataset, indicating that its dominant transcriptomic structure is preserved. Together, these results suggest that regional vulnerability collapses onto a low-dimensional molecular and cellular axis defined by rise-and-fall dynamics.

## Introduction

Neurodegenerative diseases are characterized by the accumulation of misfolded proteins in the brain, such as tau in Alzheimer’s disease and *α*-synuclein in Parkinson’s disease (PD) [1]. These proteins typically have physiological functions but contribute to neuronal dysfunction and degeneration when misfolded. Pathology does not develop uniformly across the brain. Instead, it follows reproducible spatial progression patterns in which some regions are affected earlier and pathology subsequently appears in a subset of anatomically connected regions [2–4]. These stereotyped patterns suggest that disease spreads through the brain rather than emerging independently across regions.

A leading hypothesis proposes that pathological proteins propagate in a *prion-like* manner, transferring between neurons and forming aggregates in recipient cells [5, 6]. Experimental studies support the view that misfolded *α*-synuclein can spread through connected neural circuits [7–9]. However, connectivity alone does not determine where pathology accumulates [10, 11]. Transcriptional variability, cell type composition, and other biological factors also shape the spatial distribution of pathology [8, 12]. A central challenge is therefore to understand how network-mediated propagation interacts with region-specific vulnerability to produce the observed patterns of disease.

Mathematical models have been used to address this problem. Connectome-based approaches have shown that network structure contains substantial information about the spatial distribution of pathology [8, 13–18]. More recent models incorporate local amplification dynamics and fit trajectories directly to data [19–25] and can reproduce non-monotonic pathology trajectories over time, including phases of decline [26]. However, it remains unclear how variation in these dynamics across regions relates to underlying biological factors. This gap in knowledge limits our ability to identify the biological factors that determine regional vulnerability, since these may be reflected in the temporal dynamics of pathology rather the maximum magnitude of pathology. To identify these factors, a straightforward approach would be to build models informed by data that resolve regional trajectories over time.

Mouse seeding experiments provide such data. In particular, *α*-synuclein preformed fibril (PFF) models enable quantification of pathology across hundreds of anatomically defined regions over multiple post-injection timepoints [8, 12, 16, 26–34]. This approach makes it possible to measure how pathology evolves in each region, rather than only where it accumulates. Inspection of these trajectories suggests that pathology does not follow simple monotonic accumulation over time [8, 35–38]. The biological significance of these non-monotonic pathology trajectories is not known.

Here, we analyze whole-brain histopathology measurements of *α*-synuclein in mouse models of PD from 3 days to 9 months after PFF injection into the dorsal striatum. We use a family of network dynamical system models that describe the spread of pathology across anatomical connections together with region-specific dynamics. Model classes are then compared based on their ability to reproduce the observed spatiotemporal patterns, showing that pathology burden does not simply accumulate but instead follows rise-and-fall dynamics across the brain.

To interpret the biological basis of these dynamics, we integrate whole-brain gene-expression and cell-type data with the inferred model parameters. This approach reveals a single dominant one-dimensional axis in gene expression that aligns with regional rise-and-fall dynamics, indicating that regional molecular identity is organized along a common gradient. This axis corresponds to a coordinated transcriptional program: regions with stronger rise-and-fall dynamics show increased expression of genes involved in energy metabolism, synaptic function, and protein homeostasis pathways. It is also reflected in cellular composition: regions with stronger rise-and-fall dynamics are enriched for monoamine neurons, such as dopaminergic populations. To assess generality, we generate and analyze another histopathology dataset with hippocampal PFF injection. Despite the different injection location, the same one-dimensional vulnerability axis is recovered wherein molecular and cellular associations are preserved. Together, these results suggest that rise-and-fall dynamics are a fundamental feature of *α*-synuclein pathology, and that regional vulnerability is organized along a common molecular and cellular gradient that reflects these dynamics.

## Results

### High-temporal-resolution mapping of *α*-synuclein pathology across the mouse brain

To investigate the dynamics of prion-like propagation, we need quantitative pathological measurements at a high temporal and spatial resolution. We analyzed two histopathology datasets measuring the progression of *α*-synuclein pathology across the mouse brain following focal injection of PFFs into the striatum and hippocampus, respectively [17]. Pathology burden was quantified across anatomically defined brain regions, corresponding to 412 nodes in the Allen Mouse Brain Atlas connectome [39]. Measurements were obtained at eight timepoints following injection in the striatal dataset (0.1, 0.2, 0.3, 0.5, 1, 3, 6, and 9 months) and three timepoints in the hippocampal dataset (4, 6, and 9 months) (**Fig. 1A**). The datasets are used to constrain dynamical systems models for prion-like propagation to examine the trajectories of *α*-synuclein pathology.

**Figure 1:**
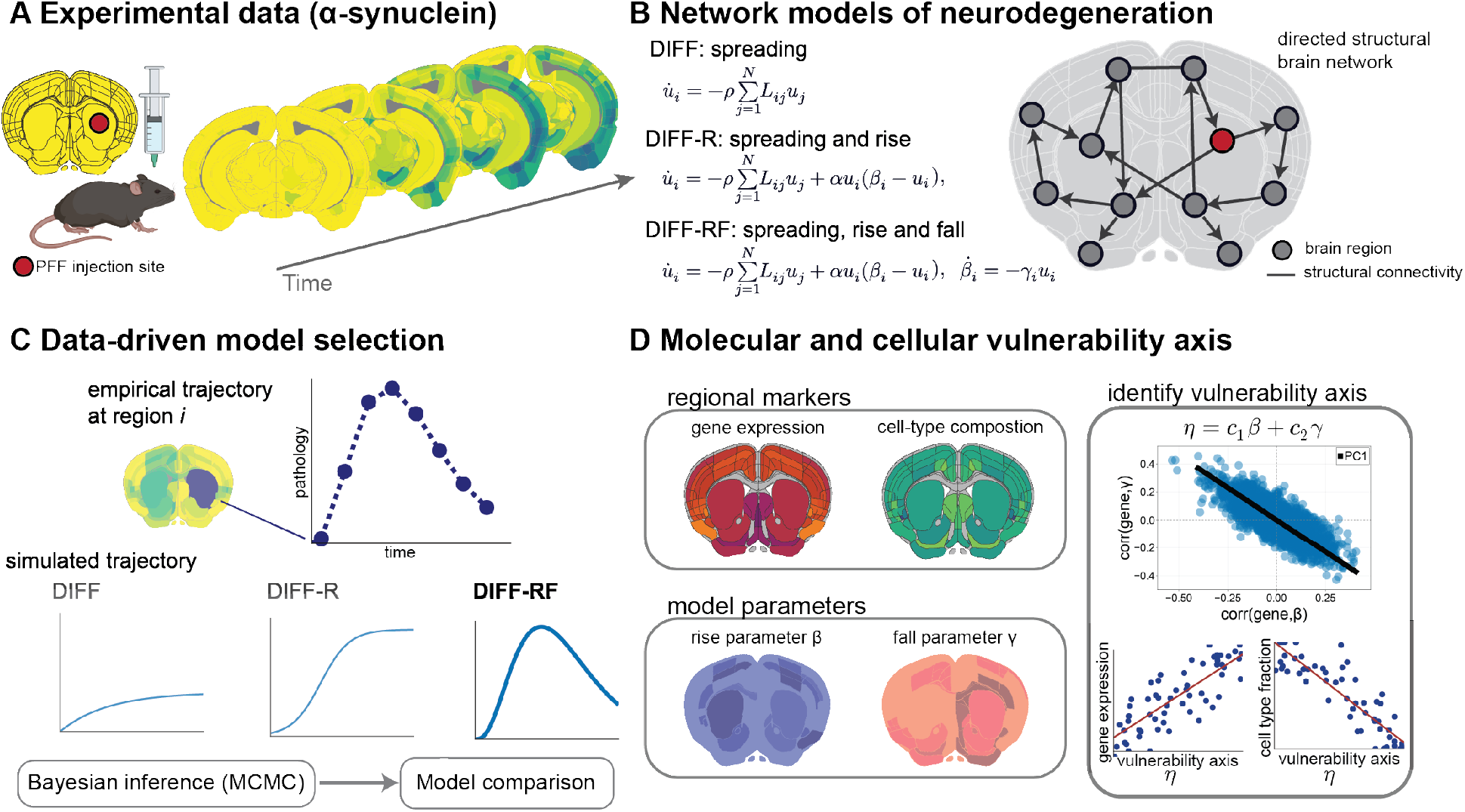
Overview of experimental data, network models, and biological interpretation. (**A**) Experimental *α*-synuclein seeding paradigm in the mouse brain, illustrating the temporal evolution of pathology following focal injection. (**B**) Network-based dynamical models of neurodegeneration. The DIFF model captures pure network spreading, DIFF-R incorporates local growth or *rise*, and DIFF-RF further includes *fall* dynamics. (**C**) Data-driven model selection. Regional empirical trajectories are compared with simulated dynamics using Bayesian MCMC inference for principled comparison of competing models. (**D**) Molecular and cellular vulnerability axis. Regional gene expression and cell-type composition are related jointly to the rise (*β*) and fall (*γ*) parameters. Principal component analysis of gene–parameter associations identifies a dominant axis *η* = *c*_1_*β* + *c*_2_*γ*, and enrichment analysis is performed along this axis to characterize associated pathways and cell types.

### Hierarchical dynamical models of *α*-synuclein propagation

To infer the dynamics of *α*-synuclein pathology, we evaluate three mathematical models for prion-like propagation against histopathological data. These models are described as systems of ordinary differential equations and form a family of network-based dynamical models that progressively increase in mechanistic complexity (**Fig. 1B**). The simplest—commonly termed the network *diffusion* model—describes passive spreading of pathology along anatomical connections and is denoted DIFF [13]. The second model augments diffusion with local aggregation dynamics governed by Fisher-Kolmogorov-Petrovsky-Piskunov (FKPP)-type growth, allowing pathology to increase within regions; this formulation has been widely used in prior work, and we denote it DIFF-R, where the “R” indicates *rise* [19]. The third model further incorporates a process that reduces regional carrying capacity in response to pathology, enabling rise-and-fall dynamics; we denote this model DIFF-RF, where the “F” indicates *fall*.

These models require a network on which the spreading occurs, a choice of modeling parameters, and initial conditions for the system variables. In this study, we use a known structural brain network, whereas we infer the modeling parameters and initial conditions. Each model inputs a directed structural connectome of the mouse brain [39], which dictates the network or graph along which the prion-like spreading occurs. We also distinguish between *global* and *regional* model parameters, where the former are constant across all regions and the latter vary region-per-region. The DIFF model has one global parameter, *ρ*, designating the time scale of spreading, and no regional parameters. The DIFF-R model has two global parameters, *ρ* and *α*, where the latter designates the time scale of regional growth, and one regional parameter, *β*_*i*_ ∈ ℝ, which we refer to as the rise parameter. It determines whether pathology grows locally and its saturation level (the carrying capacity). The DIFF-RF model has two global parameters, again *ρ* and *α*, and two regional parameters *β*_*i*_ and *γ*_*i*_. It inherits the rise parameter from DIFF-R and introduces the fall parameter *γ*_*i*_ *>* 0, which controls how rapidly pathology reduces this carrying capacity. While we use a known structural brain network, the modeling parameters and initial conditions are inferred with a Bayesian aproach as described next.

Model parameters and initial conditions were inferred using Bayesian inference with Markov chain Monte Carlo (MCMC), and model classes were compared using likelihood-based information criteria and predictive accuracy metrics, such as the Widely Applicable Information Criterion (WAIC) to select the model that best recapitulates the pathology data (**Fig. 1C**) [40]. The posterior of the modeling parameters relating to the rise (*β*_*i*_) and fall (*γ*_*i*_) were then compared with transcriptomic and cell-type whole-brain data to identify which biological factors associate with the rise and fall dynamics (**Fig. 1D**). Details of the mathematical models, parameter descriptions, parameter priors, parameter estimation procedures, and transcriptomic and cell-type data are provided in the Materials and Methods. Before comparing the three mathematical models to the pathology data, we note that the empirical connectome is directed so we evaluate which spreading direction best captures the pathology data while also comparing it to a spatial null network.

### Retrograde transport best explains *α*-synuclein propagation across the brain

Prion-like propagation may occur in the retrograde direction (from post-synaptic to pre-synaptic cell), in the anterograde direction (from pre-synaptic to post-synaptic cell), or in both directions. We examined which transport mechanism best explains the observed patterns of protein spread and compared these models against a null model in which spreading depends on physical distance between brain regions rather than propagation along anatomical connections. We considered four candidate mechanisms: retrograde transport, anterograde transport, bidirectional transport (combining retrograde and anterograde), and spatial transport independent of anatomical connectivity (which we dub Euclidean transport). We tested these mechanisms by modifying the network transport term in our mathematical models, represented by the graph Laplacian, and compared model performance using the WAIC.

Across all three models, retrograde and bidirectional transport consistently outperform anterograde and Euclidean transport (**Fig. 2**). Retrograde and bidirectional transport perform similarly across the models, with bidirectional transport narrowly outperforming retrograde transport in the DIFF model. In the DIFF-R and DIFF-RF models, retrograde and bidirectional transport are statistically indistinguishable. These results indicate that *α*-synuclein propagation is best explained by transport along anatomical connections, with a dominant retrograde component. As retrograde performs similarily to bidirectional transport and is a simpler mechanism, we continue using a retrograde transport mechanism in all subsequent computational modeling and analyses.

**Figure 2:**
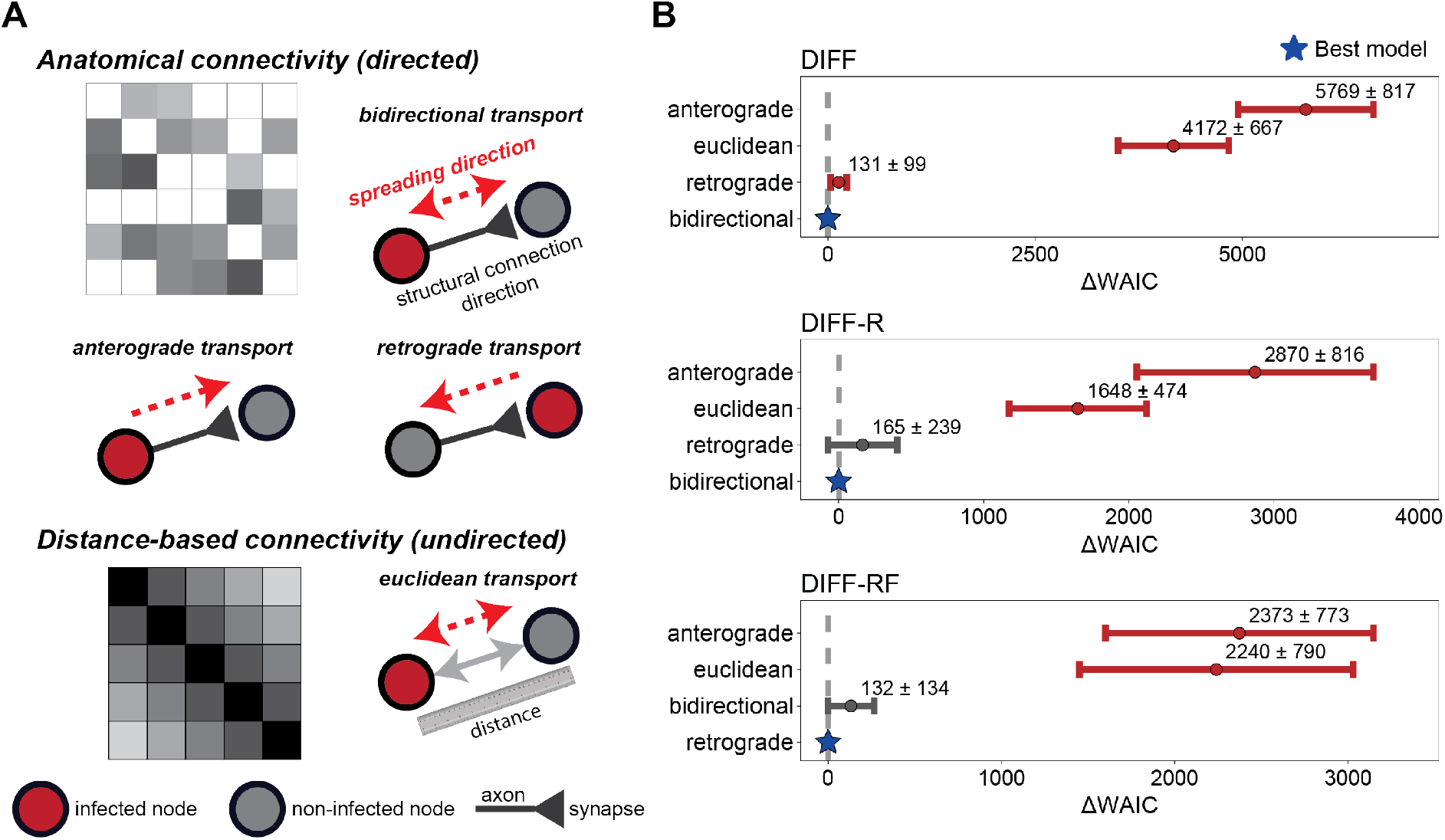
Transport mechanism hypothesis testing. (**A**) Schematic illustration of the weight matrices underlying the transport mechanisms. Anatomical transport operates on a directed structural connectivity matrix, with spreading constrained to existing axonal projections (retrograde, anterograde, or bidirectional). Euclidean transport instead uses a symmetric, distance-based weight matrix independent of anatomical connectivity. (**B**) ΔWAIC (mean ± SD across data points) for four candidate transport mechanisms—retrograde, anterograde, bidirectional, and spatial (Euclidean)—within the DIFF, DIFF-R, and DIFF-RF model classes. Lower values indicate better model performance; stars denote the best-performing mechanism within each class (ΔWAIC = 0, dashed line). Grey lines indicate that the WAIC distribution overlaps with the best performing model.

### Prion-like propagation is best captured by rise-and-fall dynamics

We next asked what dynamical process best describes the spatiotemporal trajectories of *α*-synuclein pathology. To address this question, we compared the three hierarchical model classes introduced above. By comparing these models, we test which scenarios best describe *α*-synuclein propagation: (i) propagation along structural connections, (ii) propagation along structural connections with subsequent rise in pathology, and (iii) propagation along structural connections with subsequent rise and then fall in pathology. These scenarios are represented by the DIFF, DIFF-R, and DIFF-RF models, respectively.

Our histopathological data strongly support the proposed rise-and-fall dynamics. As shown in **Fig. 3A**, DIFF-RF achieves the lowest WAIC, followed by DIFF-R and DIFF. This ordering is consistent across additional metrics (**Table 1**): DIFF-RF yields the lowest Akaike information criterion and mean squared error, whereas DIFF-R attains the lowest Bayesian information criterion, likely due to its smaller parameter dimension. We next examined predictive agreement between model-predicted and observed pathology using *maximum a posteriori* parameter estimates and *R*^2^ score based on squared deviations from the identity line, normalized by the variance of the observed values (see Materials and Methods). When pooling all regions and timepoints (**Fig. 3B**), predictive accuracy increases substantially across the model hierarchy, with *R*^2^ = 0.34 for DIFF, 0.75 for DIFF-R, and 0.84 for DIFF-RF. These differences are particularly evident in regional trajectory comparisons (**Fig. 3C**). DIFF produces slow monotonic increases and underestimates peak pathology levels. DIFF-R captures the rapid initial rise and regional saturation but fails to reproduce the subsequent decline observed in several high-burden regions. In contrast, DIFF-RF captures both the rise and the later reduction in pathology where present, while maintaining close agreement in regions that show weaker decline.

**Table 1:** Model comparison metrics. All Δ values are computed with respect to the best-performing model for each criterion (minimum value). Lower values indicate better performance.

| Model | ΔWAIC | ΔAIC | ΔBIC | ΔMSE |
| --- | --- | --- | --- | --- |
| DIFF | 18048 | 13936 | 9395 | $3.22 \times 10^{-3}$ |
| DIFF-R | 2961 | 1326 | 0 | $3.20 \times 10^{-4}$ |
| DIFF-RF | 0 | 0 | 1880 | 0 |

**Figure 3:**
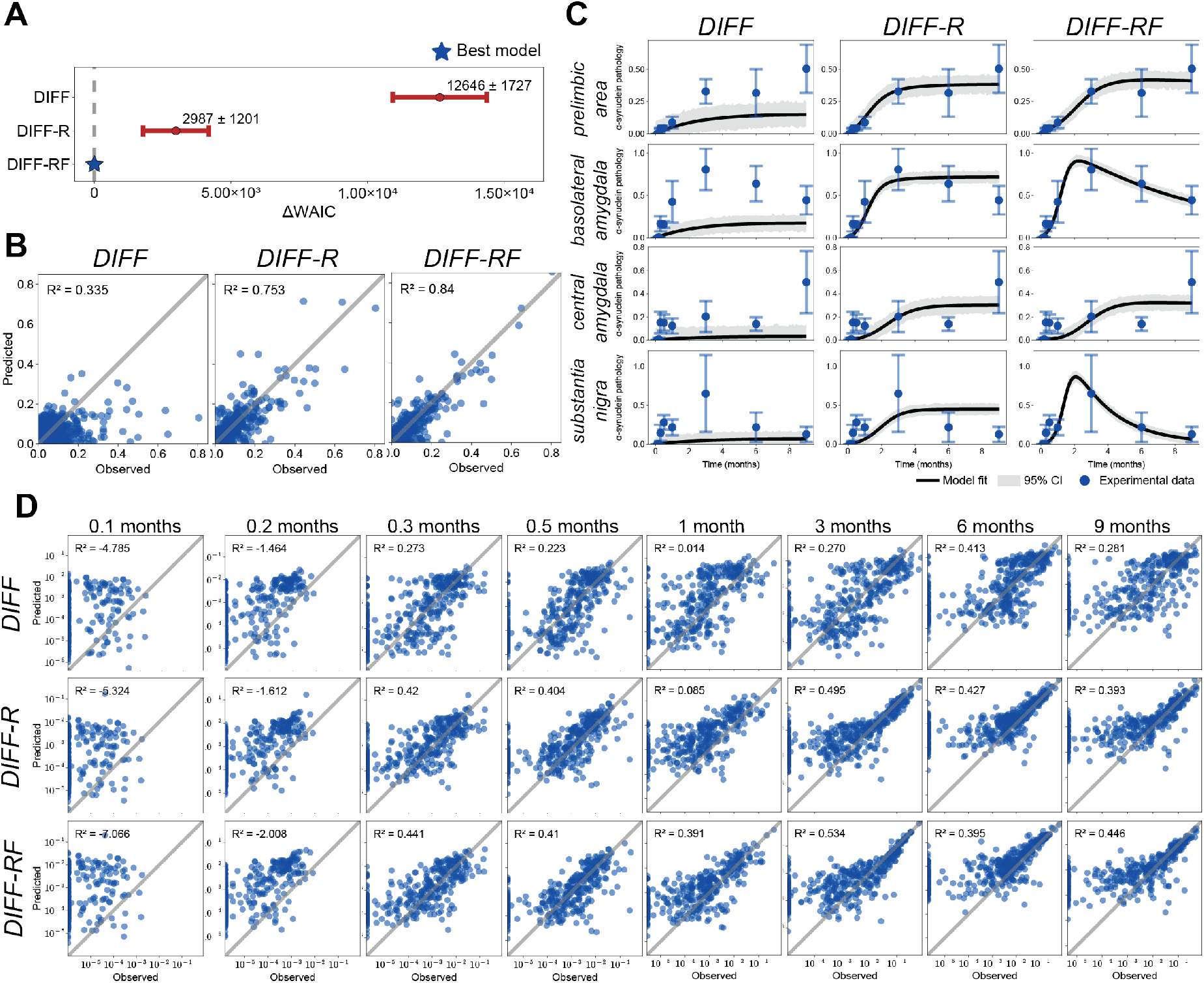
Model comparison and predictive performance across time. (**A**) Model comparison using the WAIC for DIFF, DIFF-R, and DIFF-RF. Points show mean *±* standard deviation across data samples, with values reported relative to the best-performing model (ΔWAIC = 0, dashed line). (**B**) Predicted versus observed pathology for each model, pooling all regions and timepoints. Each point corresponds to a region–timepoint pair; the diagonal indicates identity. (**C**) Regional trajectory fits for the four highest-burden regions. Black lines denote model predictions with shaded bands indicating 95% credible intervals; blue points and error bars show empirical mean and standard deviation. (**D**) Log-log predicted versus observed pathology at individual timepoints (all regions pooled). Panels correspond to increasing time after injection (0.1 to 9 months). The diagonal indicates identity.

The advantage of DIFF-RF is also evident when examining predictions at individual timepoints (**Fig. 3D**). To capture broad, overall agreement with data across time, we examine the log-log agreement between observations and model predictions. All models show weak agreement at the very earliest time points at 3 and 6 days post injection, but improve rapidly from 9 days onward. DIFF shows weak agreement at early stages and moderate agreement across the time course. DIFF-R and DIFF-RF substantially improve predictive performance, with DIFF-RF consistently reaching higher *R*^2^ across all time points, excluding at 6 months where DIFF and DIFF-R perform similarily. Comparisons in identity (not log-transformed) space give a similar picture, where DIFF-RF consistently outperforms the other models at individual time points (**Fig. S1**).

Together, these results indicate that pathology does not simply propagate and accumulate across the brain but instead follows rise-and-fall trajectories in many regions. To evaluate the robustness of these results, we next investigated the impact of the structural connectome network and the choice of injection site on model performance.

### Connectome organization and injection site influence rise-and-fall dynamics

To determine whether rise-and-fall dynamics depend on the organization of the empirical connectome and the spatial location of the simulated PFF injection site, we compared the DIFF-RF model using the empirical connectivity and injection region with variants in which either the edge arrangement or the injection location was randomized. Model performance was highest when using the empirical connectome and injection site (**Fig. 4**). Randomizing either the network topology or the seeding location consistently reduced agreement between model predictions and the observed pathology trajectories. These results indicate that accurate reproduction of the data depends on both the organization of the connectome and the anatomical origin of pathology.

**Figure 4:**
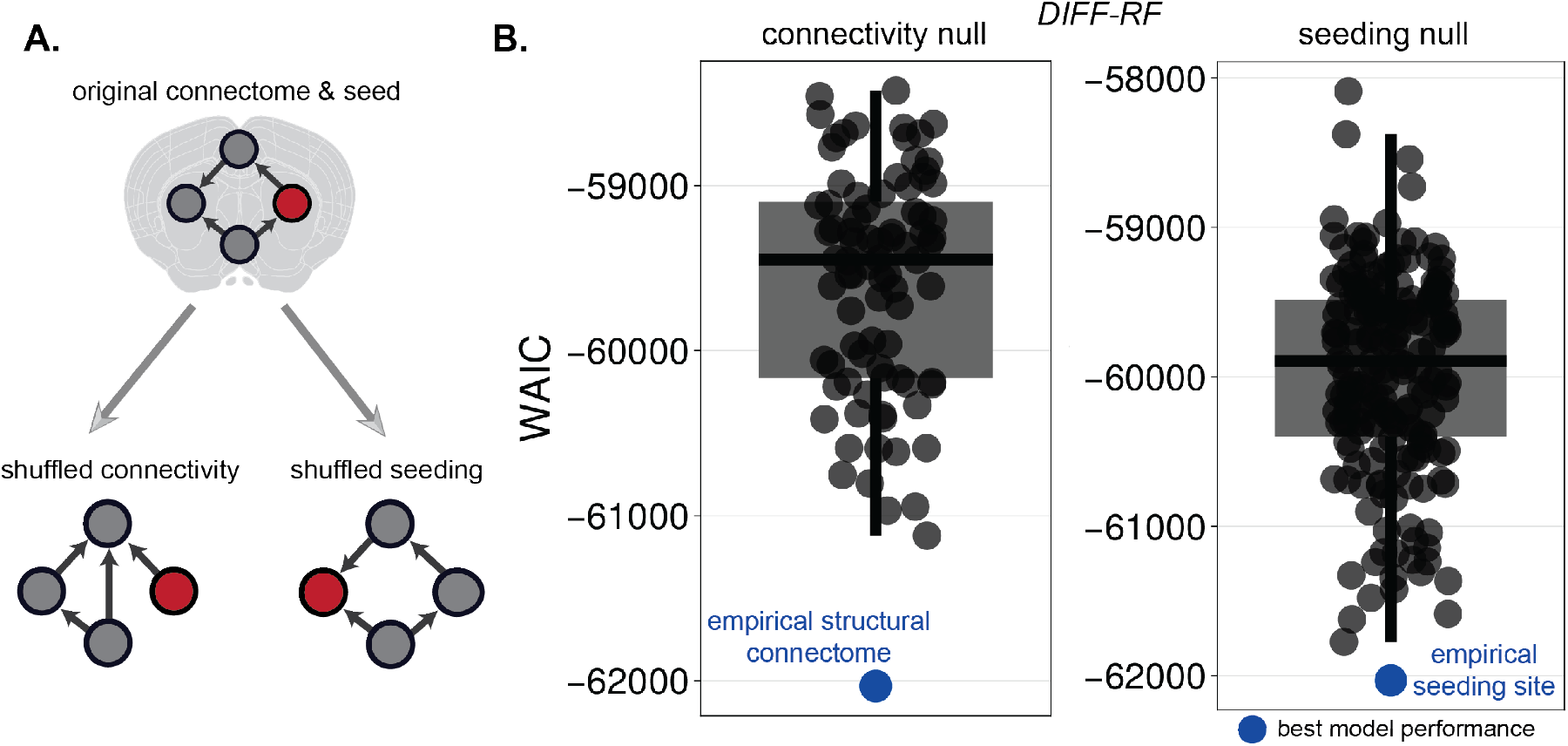
Null-model analysis of connectome organization and seeding location. (**A**) Schematic illustration of the empirical configuration and null-model perturbations. The empirical model uses the anatomical connectome and experimental seeding location. Two null variants are constructed by randomizing either the edge arrangement of the connectivity matrix (connectivity null) or the seeding location (seeding null), while keeping all other components unchanged. (**B**) WAIC values for the null models. Grey points show WAIC across randomized realizations; boxes indicate their median and interquartile range. The blue point denotes the empirical configuration.

### Rise-and-fall dynamics improve out-of-sample prediction

The rise-and-fall dynamical model is more complex and has more tunable parameters than its counterparts. To assess the robustness of the rise-and-fall advantage, we compare the out-of-sample predictive performance of the DIFF-RF and DIFF-R models. We performed a leave-one-timepoint-out analysis in which the final timepoint (*T*) was excluded from model fitting and used for evaluation. Models were fit to the remaining timepoints, and predictions were generated for the held-out observations.

Rise-and-fall dynamics improve out-of-sample predictive power. In particular, DIFF-RF shows higher predicted– observed agreement than DIFF-R (**Fig. 5A**, left) in both in-sample and out-of-sample datapoints. When evaluated on the held-out timepoint, predictive performance decreases for both models, but DIFF-RF retains substantially higher explanatory power (*R*^2^ = 0.479) than DIFF-R (*R*^2^ = 0.154) (**Fig. 5A**, right). Although one-step-ahead prediction is challenging in this dataset, the advantage of incorporating region-specific fall dynamics persists out of sample. This improvement is also evident in regional trajectory comparisons (**Fig. 5B**). DIFF-RF more closely matches the held-out pathology values in some high-burden regions, whereas DIFF-R tends to over- or under-shoot the final timepoint depending on the region. Together, these results show that incorporating rise-and-fall dynamics improves short-horizon predictive accuracy. The ability of rise-and-fall dynamics to explain and predict *α*-synuclein propagation raises the possiblity that these dynamics reflect underlying biological processes.

**Figure 5:**
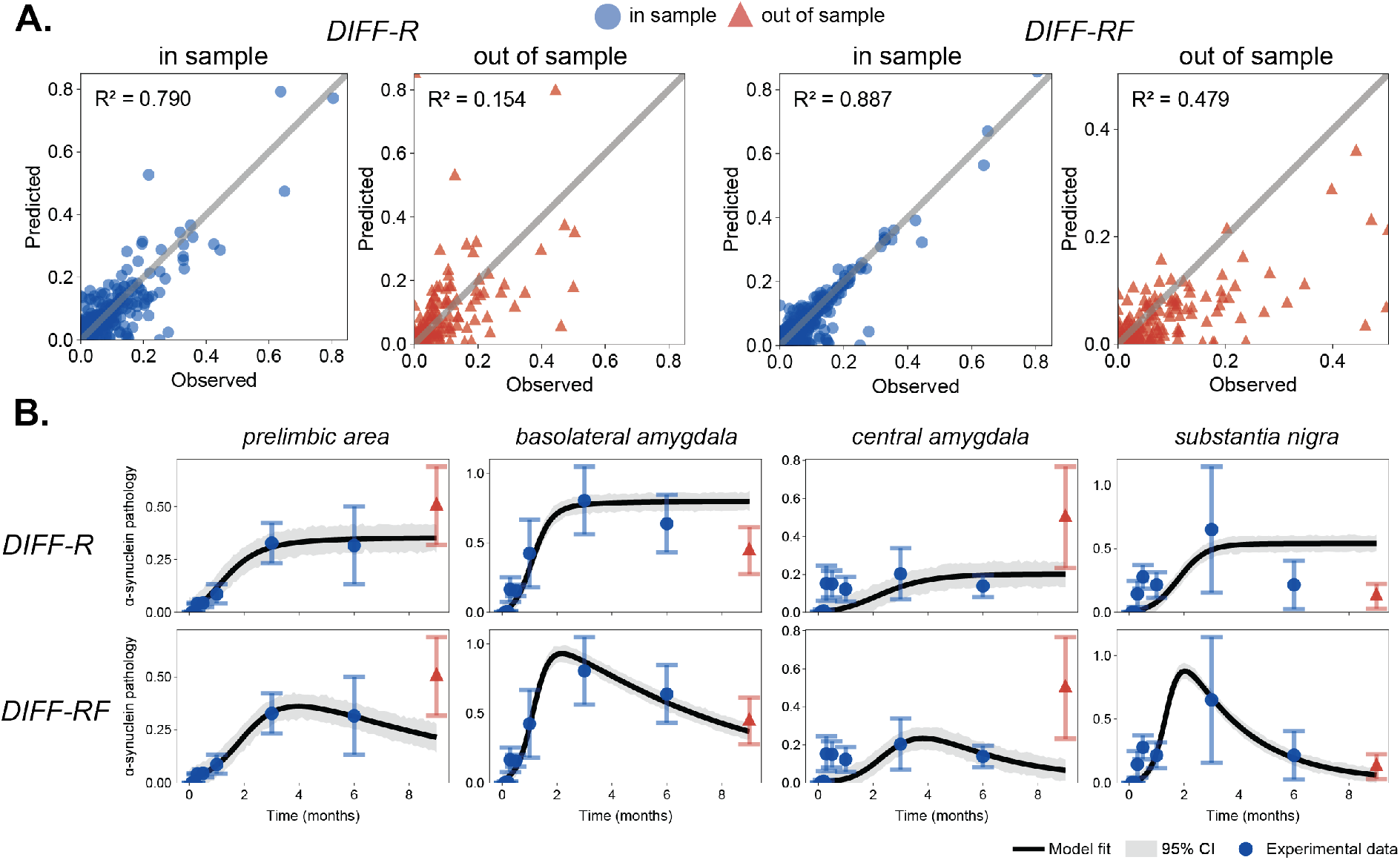
Short-horizon out-of-sample evaluation. (**A**) Predicted versus observed regional pathology for DIFF-R and DIFF-RF. Left panels show agreement across all timepoints, including the held-out observations; right panels show performance on the held-out final timepoint (*T −* 1) only. Red triangles denote held-out observations. DIFF-RF achieves higher *R*^2^ both overall and on the held-out timepoint. (**B**) Regional trajectories for the four highest-burden regions, with the final timepoint excluded from fitting. Black lines show model predictions; blue circles denote fitted data; red triangles denote held-out observations. DIFF-RF more closely captures the held-out pathology levels than for DIFF-R.

### Inferred rise-and-fall parameters reveal a molecular and cellular vulnerability axis

To investigate whether rise-and-fall dynamics reflect underlying biological structure, we examine whether the inferred model parameters relate to gene expression. For each gene, we fit a multiple regression model relating regional expression levels to the *z*-transformed rise-and-fall parameters, denoted *z*(*β*_*i*_) and *z*(*γ*_*i*_), where each parameter was standardized across regions (see Materials & Methods). This procedure yields a pair of coefficients describing how strongly each gene’s expression varies with the two rise-and-fall parameters across brain regions. Each gene can be represented as a point in a two-dimensional coefficient space defined by its correlational associations with *z*(*β*_*i*_) and *z*(*γ*_*i*_) (**Fig. 6A**). Principal component analysis (PCA) of this coefficient space revealed a dominant axis capturing the majority of the variance across genes. The first principal component explained 95.5% of the variance and corresponded to the direction *η* = *−* 0.95 *z*(*β*) + 0.33 *z*(*γ*), indicating a joint transcriptional gradient associated with lower rise and higher fall parameters.

**Figure 6:**
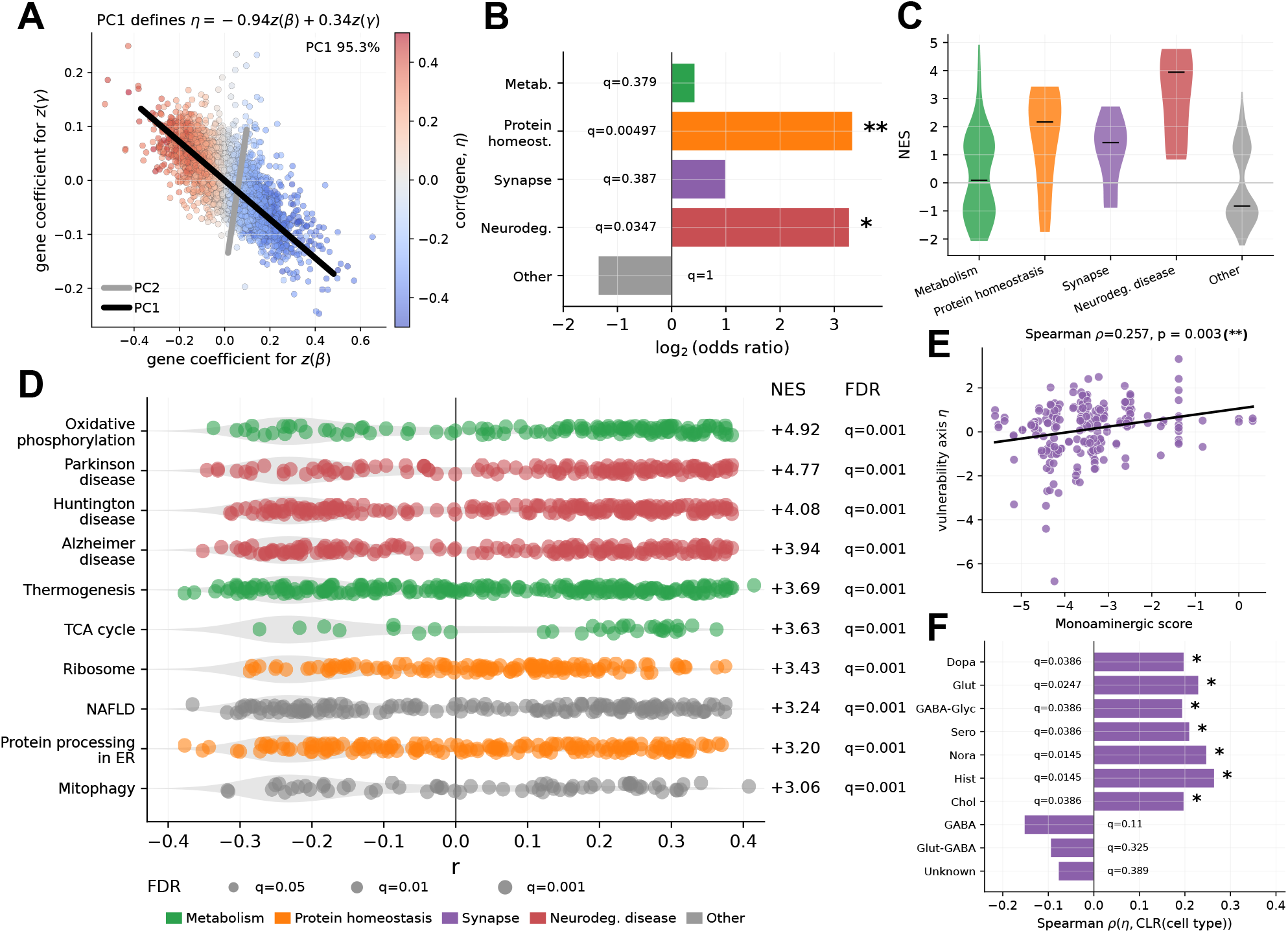
PCA-derived vulnerability axis and biological associations. (**A**) PCA of gene-level regression coefficients for rise (*β*) and fall (*γ*). Each point represents a gene in the space of coefficients for *z*-transformed rise-and-fall parameters *z*(*β*) and *z*(*γ*). Black lines denote the first and second principal components (PC1 and PC2). The first principal component defines the axis *η*. Points are colored by their Pearson correlation between gene expression and *η*. (**B**) Enrichment of functional gene categories along the vulnerability axis, shown as log_2_ odds ratios with Benjamini– Hochberg adjusted *q*-values. (**C**) Distribution of NES across functional categories. (**D**) GSEA from correlations with the vulnerability axis. Each row shows the distribution of gene-level correlations within a pathway; colored points indicate pathway membership, with normalized enrichment scores (NES) and FDR-adjusted pathway-level *q*-values shown on the right. Point size denotes the FDR-adjusted pathway *q*-value. (**E**) Relationship between vulnerability and monoaminergic cell-type scores across regions, with Spearman correlation shown. (**F**) Association between the vulnerability axis and cell-type signatures, quantified using the Spearman correlation between *η* and CLR-transformed cell-type expression.

To interpret the biological processes associated with this axis, we examine the associations between gene expression and the dominant axis *η* across regions. Each region has a value *η*_*i*_ as a function of its rise and fall parameters, where larger *η*_*i*_ indicates a stronger rise-and-fall component and that the region is high on the vulnerability axis. Investigating the correlation between the vulnerability axis *η* and gene expression across brain regions will tell us what biological pathways are associated with stronger rise-and-fall dynamics. We ranked genes by their correlation (Pearson *r*) with *η* and performed pre-ranked gene set enrichment analysis (GSEA). Genes aligned with increasing *η* were strongly enriched for pathways related to neurodegenerative disease and protein homeostasis (**Fig. 6B–D**), as well as energy metabolism pathways such as oxidative phosphorylation. Genes positively aligned with this axis (*η*) correspond to regions exhibiting stronger fall and weaker rise dynamics, indicating that proteostasis and metabolic transcriptional programs are upregulated along the direction of increasing *γ* and decreasing *β*.

We next examined whether this molecular axis was reflected in regional cellular composition. Using an independent whole-brain cell-type atlas [41], we related neurotransmitter-class fractions to the regional values of *η*. Regions with higher *η*_*i*_—corresponding to stronger fall dynamics and lower rise—showed higher fractions of monoaminergic neurons (Spearman *ρ* = 0.257, permutation *p*-value = 0.003; see Materials and Methods), with the strongest associations observed among monoaminergic neurotransmitter classes (**Fig. 6E,F**).

Together, these results indicate that the inferred parameter landscape is organized along a single molecular and cellular vulnerability axis. Regions enriched in synaptic and proteostasis transcriptional programs and populated by monoaminergic neuronal populations tend to exhibit rise-and-fall dynamics. This pattern suggests that vulnerability is associated with rise-and-fall trajectories, defining a characteristic dynamical signature of regional susceptibility. We next ask whether the vulnerability axis is preserved in a new *α*-synuclein propagation histopathological dataset.

### Molecular and cellular vulnerability axis replicates across datasets

To test whether the molecular and cellular vulnerability axis identified above generalizes across experimental conditions, we generated and analyzed a distinct whole-brain histopathology dataset in which *α*-synuclein PFFs were injected into the hippocampus instead of the dorsal striatum (**Fig. S2**). We fit the DIFF-RF model to this dataset and obtained strong agreement between predicted and observed regional pathology (*R*^2^ = 0.97 for pooled regions and timepoints; **Fig. S3**). Posterior diagnostics for the hippocampal model are shown in **Fig. S4**. We then ask whether we recover the vulnerability axis found in the striatal dataset.

We repeated the joint transcriptomic analysis relating regional gene expression to the inferred rise (*β*_*i*_) and fall (*γ*_*i*_) parameters and successfully reproduced a similar vulnerability axis. As in the striatal dataset, each gene was represented by its regression coefficients with respect to *z*(*β*_*i*_) and *z*(*γ*_*i*_), and PCA was used to summarize the dominant structure of this coefficient space. The first principal component explained 68.3% of the variance in gene coefficients and defined the axis *η* = *−* 0.97 *z*(*β*) + 0.23 *z*(*γ*), indicating a transcriptional gradient aligned with increasing fall dynamics and decreasing rise (**Fig. 7A**). Thus, despite the different PFF injection location, the dominant molecular axis again corresponds to a similar rise-and-fall dynamical direction identified in the striatal dataset.

**Figure 7:**
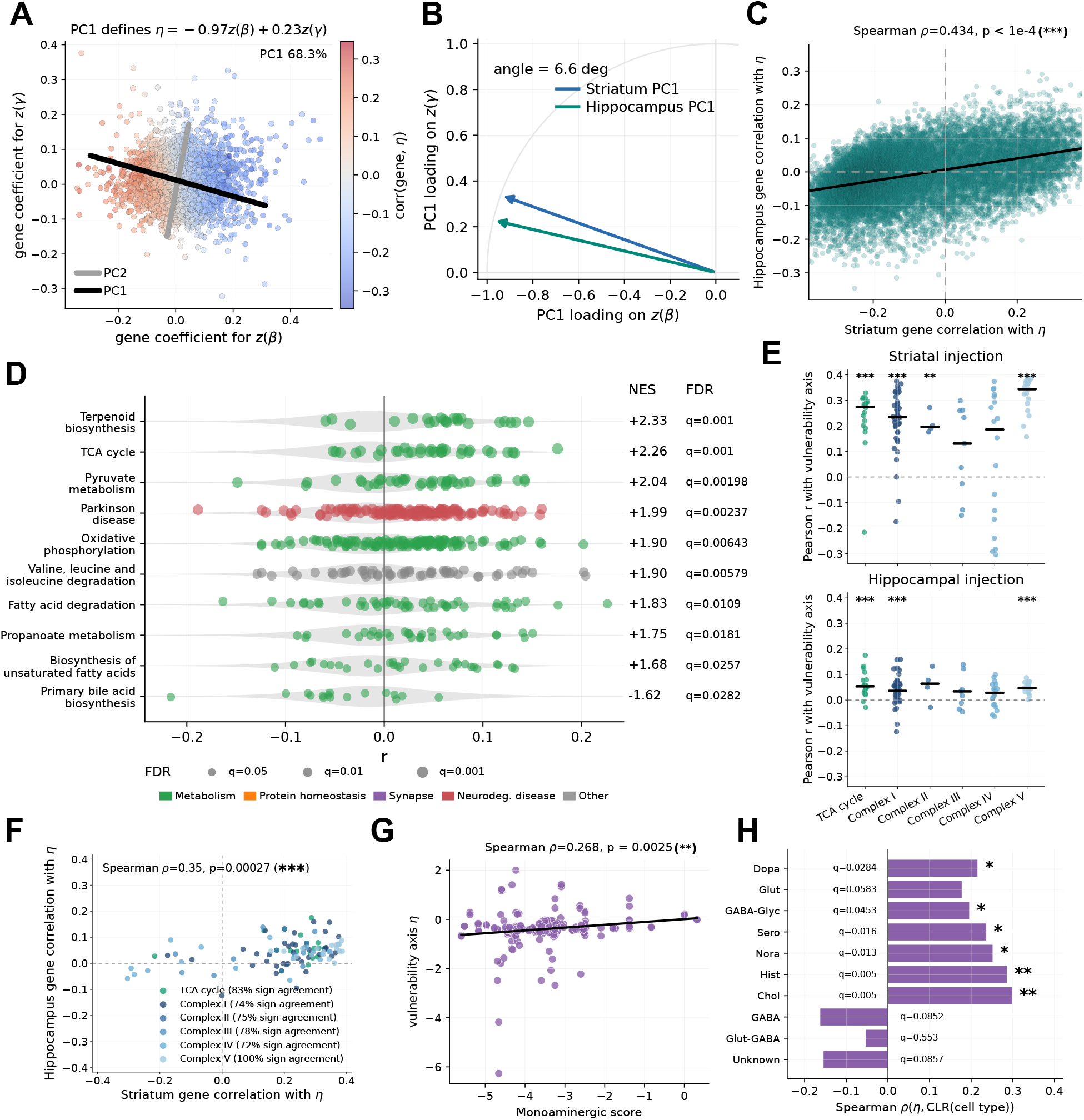
Replication of the PCA-derived vulnerability axis in hippocampal seeding. (**A**) PCA of gene-level regression coefficients for rise (*β*) and fall (*γ*) in the hippocampal dataset. Each point represents a gene; PC1 defines the vulnerability axis *η*. (**B**) Comparison of PC1 directions between striatal and hippocampal datasets. (**C**) Concordance of gene-level associations with the vulnerability axis across datasets. Each point shows the correlation of a gene with *η* in the striatal dataset (x-axis) versus the hippocampal dataset (y-axis). (**D**) GSEA for pathways significantly enriched in both striatal and hippocampal analyses, ranked by hippocampal NES. Each row shows the distribution of gene-level correlations within a pathway in the hippocampal dataset; colored points indicate pathway membership, with NES and FDR-adjusted *q*-values shown on the right. (**E**) Gene-level correlations with the vulnerability axis for curated TCA-cycle and oxidative phosphorylation genes in the striatal and hippocampal datasets. Stars denote one-sided one-sample *t*-tests for positive mean correlation within each pathway module. (**F**) Concordance of curated energy-metabolism gene correlations with *η* across datasets. (**G**) Relationship between vulnerability and monoaminergic cell-type scores across regions. (**H**) Association between the vulnerability axis and cell-type signatures, quantified using Spearman correlation between *η* and CLR-transformed cell-type expression.

To directly compare the two experiments, we next examined the alignment of the vulnerability axes across the datasets. In the standardized (*z*(*β*_*i*_), *z*(*γ*_*i*_)) coefficient space, the first principal components from the striatal and hippocampal analyses are highly similar (cosine similarity = 0.993, angle = 6.6^*°*^; **Fig. 7B**), indicating near-identical vulnerability axes. In the raw (*β, γ*) coefficient space, the first principal components were also highly aligned (**Fig. S5**). At the level of individual genes, correlations with the axis *η* were moderately preserved across datasets (Spearman *ρ* = 0.434, *p <* 10^*−*4^; **Fig. 7C**), indicating that the dominant transcriptomic gradient is reproducible across injection sites.

Pre-ranked GSEA along this axis reproduced the same major biological themes observed previously, with the strongest overlap in metabolic pathways. Among pathways significantly enriched in both datasets, the largest hippocampal enrichment scores included terpenoid backbone biosynthesis (NES = 2.33, *q* = 0.001), citrate cycle (NES = 2.26, *q* = 0.001), pyruvate metabolism (NES = 2.04, *q* = 0.002), Parkinson disease (NES = 1.99, *q* = 0.002), and oxidative phosphorylation (NES = 1.90, *q* = 0.006; **Fig. 7D, Fig. S6**). At the category level, the hippocampal axis was enriched for metabolism (log_2_ odds ratio = 2.74, *q* = 0.0035), and neurodegenerative disease pathways (log_2_ odds ratio = 3.98, *q* = 0.049). Thus, the hippocampal replication recapitulates the energy-metabolic and Parkinson-related transcriptional programs identified in the striatal experiment.

Because metabolism was the most consistent enriched category across datasets, we next asked whether core energy-metabolism is elevated or lowered along the vulnerability axis. We curated genes encoding core TCA-cycle enzymes and oxidative phosphorylation complexes (**Table S1**) and examined their correlations with *η*. Gene correlations were predominantly positive in both datasets, indicating that higher expression of energy-metabolism is associated with increasing vulnerability-axis values (**Fig. 7E**). Using one-sided one-sample t-tests (see Materials and Methods), this pattern was significant for the TCA cycle in both the striatal dataset (*t*(17) = 7.91, *p* = 2.1 × 10^*−*7^) and hippocampal dataset (*t*(17) = 5.13, *p* = 4.2 × 10^*−*5^), and was also significant in both datasets for Complex I (striatum: *t*(37) = 11.07, *p* = 1.3 *×* 10^*−*13^; hippocampus: *t*(37) = 3.99, *p* = 1.5 *×* 10^*−*4^) and Complex V (striatum: *t*(16) = 19.10, *p* = 9.7 *×* 10^*−*13^; hippocampus: *t*(16) = 9.20, *p* = 4.3 *×* 10^*−*8^). Gene-level correlations for these curated energy-metabolism genes were also preserved across datasets (Spearman *ρ* = 0.350, *p* = 2.7 *×* 10^*−*4^; **Fig. 7F**). These results indicate elevated energy metabolism along the vulnerability axis.

The cell-type analysis in the hippocampal data yielded a similar result to the striatal dataset. Regional values of *η* were again positively associated with monoaminergic neuronal composition (Spearman *ρ* = 0.268, *p* = 0.0025), and dopaminergic neurons again showed positive associations among neurotransmitter classes (**Fig. 7G,H**).

The rise parameter (*β*_*i*_) shows a stronger agreement across the datasets than the fall parameter (*γ*_*i*_). Multiple linear regression coefficients associated with *β*_*i*_ showed a moderate positive correspondence across datasets (Spearman *ρ* = 0.414, *p <* 10^*−*4^, 64% sign agreement; **Fig. S7B**), whereas coefficients associated with *γ* showed a weak negative correspondence (Spearman *ρ* = *−*0.162, *p <* 10^*−*4^, 40% sign agreement; **Fig. S7A**).

## Discussion

Our results show that *α*-synuclein pathology exhibits rise-and-fall dynamics, and that a disease-relevant vulnerability axis is encoded in these dynamics. These findings indicate that regional vulnerability is not determined solely by the capacity to accumulate pathology, but by the temporal trajectories of pathology. The dynamics of brain-wide pathology may therefore provide important insight into regional susceptibility to *α*-synuclein pathology. These findings also extend connectome-based models of prion-like propagation by showing that temporal trajectories impose additional constraints on pathology dynamics, consistent with findings in experimental and modeling studies reporting biphasic pathology trajectories [26, 35–38]. Here, we show that these dynamics are regionally structured and organized along a low-dimensional axis linked to transcriptional programs and cell-type composition.

### Biological underpinnings of fall dynamics

A plausible interpretation of the fall parameter is that it reflects neuronal loss following initial accumulation of *α*-synuclein pathology. Experimental seeding studies show that pathological *α*-synuclein induces progressive neurodegeneration, including neuritic degeneration and loss of vulnerable neuronal populations [27, 42]. In this context, a reduction in measured pathology is expected if affected neurons or their projections are lost. Network-based analyses further support this interpretation, showing that model-derived vulnerability aligns with gene programs regulating neuronal survival and that perturbation of these pathways reduces both pathology and neuron loss [8, 17]. Together, these results support interpreting fall dynamics primarily as degeneration of vulnerable neuronal populations.

However, fall dynamics are unlikely to reflect a single mechanism. Whole-brain modeling integrating gene expression identifies later phases of reduced pathology attributed to local processes beyond spreading [26]. Intracellular degradation pathways, including the autophagy–lysosomal and ubiquitin–proteasome systems, modulate *α*-synuclein levels in vivo [43, 44], and microglia contribute to extracellular aggregate clearance [45]. These processes can reduce pathology independently of neuronal loss. Fall dynamics may therefore be interpreted as a phenomenological description possibly reflecting a combination of degeneration, clearance processes, and consolidation.

### Interpreting the cellular and transcriptomic vulnerability axis

We show by transcriptional analysis that rise-and-fall dynamics are organized along a one-dimensional axis. This structure arises at the level of gene associations, not the parameters themselves. The rise and fall parameters vary independently across regions, but the genes associated with these parameters align along a common axis. Prior work has linked network-derived vulnerability to regional gene expression, identifying transcriptional correlates of susceptibility [17, 32, 33, 46, 47]. In the present results, these associations are organized along a low-dimensional axis that emerges from the joint structure of rise and fall dynamics. The vulnerability axis therefore represents a distinct organizational feature of regional molecular variation associated with propagation dynamics.

The enriched transcriptional programs provide a mechanistic interpretation of this axis. Regions with stronger fall and weaker rise dynamics are associated with genes involved in energy metabolism, protein homeostasis, and synaptic function. Mitochondrial dysfunction increases susceptibility to *α*-synuclein aggregation and toxicity [48], while disruption of proteostasis alters aggregation dynamics *in vivo* [43]. Synaptic activity modulates *α*-synuclein release and transmission, and pathological aggregation impairs synaptic function and promotes synapse loss [49, 50]. Energy metabolism pathways were especially well conserved in the vulnerability axis across datasets. Moreover, key enzymes in the TCA cycle and oxidative phosphorylation were up-regulated, indicating elevated expression of core metabolic machinery in regions with more pronounced rise-and-fall trajectories. Previous studies support this interpretation. Vulnerable nigral dopaminergic neurons have elevated basal oxidative phosphorylation, limited respiratory reserve, and increased oxidative stress, suggesting that high energetic demand can contribute to selective vulnerability [51]. Oxidative and nitrative stress can in turn promote *α*-synuclein aggregation and spreading in vivo [52– 54]. Together, these findings suggest that elevated metabolic gene expression along the vulnerability axis may reflect increased energetic demand that predisposes regions to metabolic stress and *α*-synuclein pathology.

The association with monoaminergic neuronal composition provides a complementary cellular interpretation. Regions with stronger rise-and-fall dynamics are enriched for monoaminergic neurons, particularly dopaminergic populations. This is consistent with a recent computational study finding that monoaminergic neuronal populations are strong mediators of prion-like *α*-synuclein propagation [47]. Moreover, dopaminergic neurons exhibit autonomous pacemaking and sustained calcium influx, leading to mitochondrial oxidant stress and increased vulnerability [55]. Similar vulnerability extends to other monoaminergic systems: noradrenergic neurons develop *α*-synuclein pathology and dysfunction in targeted models [56], and serotonergic neurons exhibit neuritic degeneration following *α*-synuclein expression [57]. Manipulating noradrenergic activity alters dopaminergic neuron survival in synucleinopathy models [58]. These results support interpreting the vulnerability axis as reflecting broader monoaminergic susceptibility. The replication of this axis across striatal and hippocampal datasets indicates that it reflects an underlying biological gradient. The hippocampal-derived axis preserves the same biological associations and direction in parameter space, but shows a weaker one-directional component, possibly due to its lower pathology levels and temporal resolution. Despite differences in injection site and propagation patterns, the same vulnerability axis is identified.

### Retrograde transport best describes *α*-synuclein propagation

Our results are consistent with previous findings identifying a primarily retrograde direction in prion-like *α*-synuclein propagation. Experimental and modeling studies consistently support a strong retrograde component in *α*-synuclein propagation, with connectivity-based predictions improving when retrograde projections are emphasized [8, 9, 30]. While anterograde spread may occur, the overall pattern across studies is consistent with retrograde-dominated transport. The rise-and-fall dynamical model provides further support for this hypothesis.

### Methodological considerations

Several limitations should be noted. We assume a fixed structural connectome which does not account for connectivity changes caused by pathology. Moreover, the model is deterministic and does not capture the increase in variance observed in histopathological samples at later timepoints. This could be achieved by modeling the prion-like propagation as a stochastic differential equation system [15], though computationally this may be prohibitively expensive with a Bayesian MCCM approach for parameter estimation. In future studies, rise-and-fall dynamics may also incorporate regional cell-type composition into the parameterization of each region, capturing the differential response to pathology as has been done in previous studies [47]. Finally, the rise-and-fall dynamical model does not incorporate neuronal activity, despite evidence that activity-dependent processes influence *α*-synuclein propagation and that coupling activity to spreading can alter system dynamics [59–62].

## Conclusion

We identify rise-and-fall dynamics as a fundamental feature of *α*-synuclein propagation and show that regional vulnerability is encoded in these dynamics, revealing a dominant and reproducible vulnerability axis linked to monoaminergic neuronal composition, energy metabolism and proteostatic transcriptional programs.

## Materials and Methods

### Striatal seeding pathology dataset

We analyzed previously published whole-brain histopathology data from mouse models of *α*-synuclein propagation following unilateral striatal injection of *α*-synuclein PFFs [17]. Pathology burden was quantified using pS129 *α*-synuclein immunostaining and registered to the Allen Mouse Brain Atlas CCFv3, yielding region-wise measurements across anatomically defined brain regions. In the present study, we used the regional pathology data aggregated to the brain regions included in the connectome model. Measurements were available at eight post-injection timepoints (0.1, 0.2, 0.3, 0.5, 1, 3, 6, and 9 months). Details of animal procedures, tissue processing, staining, image segmentation, atlas registration, and pathology quantification are provided in the original publication [17].

### Hippocampal seeding pathology dataset

#### Immunohistochemistry

Tissue was obtained from previously described mice [63]. All slides were deparaffinized with two sequential xylene baths (5 min each) and then incubated for 1 minute in a descending series of ethanol baths: 100%, 100%, 95%, 80%, 70%. After a rinse in distilled water, antigen retrieval was performed using citric acid (pH = 6; Vector Laboratories, Cat# H-3300) for 15 min at 95 °C. Slides were allowed to cool for 20 min at room temperature and washed in running tap water for 10 min.

To quench endogenous peroxidase activity, slides were incubated in 7.5% hydrogen peroxide for 30 min at room temperature. Slides were washed for 10 min in running tap water, placed for 5 min in 0.1 M Tris buffer (pH = 6), and then blocked for 1 hour at room temperature in 0.1 M Tris with 2% fetal bovine serum (FBS). Slides were incubated overnight at 4 °C in primary antibody diluted in 0.1 M Tris/2% FBS in a humidified chamber.

The following primary antibody was used: rabbit polyclonal anti-pS129 *α*-synuclein (Abcam, Cat# ab51253, RRID:AB 869973). Primary antibodies were rinsed off with 0.1 M Tris for 5 min and then incubated with the appropriate secondary antibody: goat anti-rabbit biotinylated IgG (1:1000; Vector, Cat# BA1000) in 0.1 M Tris/2% FBS for 1 hour at room temperature.

Slides were rinsed using 0.1 M Tris for 5 min, then incubated with avidin–biotin solution (Vector, Cat# PK-6100) for 1 hour. Slides were then rinsed for 5 min with 0.1 M Tris and developed with ImmPACT DAB peroxidase substrate (Vector, Cat# SK-4105). Slides were counterstained for 15 seconds with Harris hematoxylin (Fisher, Cat# 67-650-01). Slides were washed in running tap water for 5 min, dehydrated in ascending ethanol baths (70%, 80%, 95%, 100%, 100%; 1 min each), and incubated in two sequential xylene baths (5 min each). Slides were mounted with coverslips using Cytoseal mounting media (Fisher, Cat# 23-244-256). Slides were scanned at 20*×* magnification using an Aperio ScanScope XT. Digitized images were used for quantitative pathology.

#### Mouse Brain Segmentation and Registration

##### Segmentation

Scanned slides were imported into QuPath v0.5.0 (RRID: SCR 018257) [64] for analysis. A pixel classifier thresholding fluorescent intensity for the pS129 *α*-synuclein channel was applied to each section to detect positive pathology signal. Signal intensity was optimized by adjusting min/max display settings, and a consistent threshold was applied across cohorts.

##### Mouse Coronal Section Brain Registration

Images were registered to the Allen Brain Atlas CCFv3 using a modified QUINT workflow (RRID: SCR 023856) [65]. RGB images of each section were exported from QuPath as PNG files and downsampled by a factor of 12 for spatial registration. Segmentation images were generated by exporting color-coded classified pixels or cells on a white background for use in Nutil (RRID: SCR 017183) [66].

Brain images were uploaded to DeepSlice (RRID: SCR 023854) [67], where a deep neural network was used to automatically align sections to atlas regions. Preliminary alignment was performed in QuickNII (RRID: SCR 016854) [68]. Following registration, a JSON file was exported for use in VisuAlign (RRID: SCR 017978).

In VisuAlign, brain sections were refined using nonlinear transformations. Anchor points were placed on the atlas overlay and adjusted to corresponding locations in the tissue sections. Outer contours were aligned first, followed by refinement of internal structures. Final alignments were exported as FLAT and PNG files for use in Nutil.

Nutil was used for quantification and spatial analysis of classified cell types across brain regions. Each segmentation was processed using input masks from QuPath, JSON anchoring files from QuickNII, and transformation files from VisuAlign. Outputs included object area, region area, and area occupied for each classification within regions of the Allen Brain Atlas CCFv3. These outputs were used for downstream modeling and visualization.

#### Anatomical Analysis Heatmaps

##### Nutil-to-Usable

Nutil-to-Usable (N2U) is an R-based Shiny application developed at the Van Andel Institute for generating anatomical heatmaps based on the Allen Brain Atlas (RRID: SCR 024753). Outputs from Nutil were uploaded separately for left and right hemispheres.

An annotation file was generated to label metadata for each section (mouse ID, sex, treatment, post-injection interval, etc.). Checkpoint files combining raw Nutil outputs and metadata were used to accelerate downstream processing. Settings were defined to specify the level of summary and variable of interest. “Load”, defined as the ratio of pathology area to region area, was used as the primary variable when generating heatmaps of total pathology and cell body inclusion area.

##### Mouse Brain Heatmaps (MBH)

Plotting matrices were uploaded to MBH to generate anatomical heatmaps. MBH plots the compiled data for the parameter of interest represented in the plotting matrix set by N2U. Applicable coronal section figures were listed in the app by number based on the ordering of Allen Brain Atlas sections. Regional Resolution for the plots was set to either daughter (by layer) or parent (compiling layers). Plots were downloaded as svgs.

### Structural connectivity

Structural connectivity was obtained from the Allen Mouse Brain Connectivity Atlas as described previously [8, 17]. We used the directed projection matrix defined on *N* = 412 Allen CCFv3 brain regions, where each element *W*_*ij*_ represents the strength of axonal projections from region *j* to region *i* (see more details in the Mathematical Modeling Section).

To assess whether model performance depends specifically on anatomical connectivity, we also considered a spatial null model based on Euclidean distance between brain regions. In this control model, the connectivity matrix was replaced by a symmetric weight matrix constructed from inter-regional Euclidean distances between region centroids, following the procedure used in previous work [17]. This distance-based matrix preserves the spatial embedding of brain regions while removing anatomical connectivity structure.

### Gene expression data

Regional gene-expression data were obtained from the mouse spatial transcriptomic dataset from previous work [17]. Expression profiles were provided for a subset of brain regions defined in the Allen Mouse Brain Atlas CCFv3. The dataset includes measurements for 132 regions in a single hemisphere.

To align transcriptomic data with hemisphere-specific model parameters, gene-expression values were duplicated across hemispheres, assigning identical expression profiles to the left and right instances of each region. This enabled direct comparison with hemisphere-specific parameter estimates without averaging across hemispheres.

All gene–parameter analyses were performed at the hemisphere level, after matching regions between the transcriptomic dataset and the connectome model.

### Cell-type composition data

Regional cell-type composition was estimated using the Allen Whole Mouse Brain (WMB) cell taxonomy [41]. This dataset defines transcriptomic cell clusters across the mouse brain using single-cell RNA sequencing together with spatial transcriptomic mapping to the Allen Common Coordinate Framework (CCFv3) [41]. Each cluster is annotated by neurotransmitter class and associated with spatial frequency maps across anatomical regions.

Cluster sizes were obtained from the reported number of cells assigned to each transcriptomic cluster. Regional cell counts were then estimated by distributing the cells of each cluster across brain regions according to the reported spatial frequencies. These estimated counts were aggregated across clusters belonging to the same neurotransmitter class to obtain regional counts for each cell type.

When spatial frequencies were reported for anatomical structures that did not exactly match the connectome regions, contributions were propagated through the Allen brain ontology so that descendant regions contributed to their parent structures. This procedure produced estimated cell counts for every connectome region included in the model.

Regional cell-type fractions were computed by normalizing the estimated counts of each neurotransmitter class by the total estimated number of cells in that region. These fractions were used as measures of regional cell-type composition in subsequent analyses.

### Mathematical modeling

#### DIFF: Spreading-only

Let *W* ∈ ℝ ^*N ×N*^ denote the directed weighted adjacency matrix of the mouse structural connectome, where *W*_*ij*_ represents the strength of the projection from region *j* to region *i*. We then define the out-degree graph Laplacian

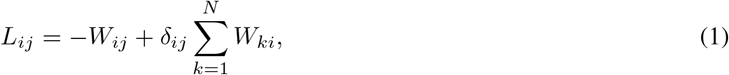

where *δ*_*ij*_ is the Kronecker delta and Σ _*k*_ *W*_*ki*_ corresponds to the total outgoing projection strength from region *i*. The spreading-only (linear diffusion) model is given by

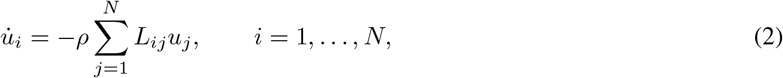

where *u*_*i*_(*t*) denotes pathology burden in region *i* and *ρ >* 0 is the transport timescale.

For anterograde transport, the directed connectivity matrix was used as defined above. For retrograde transport, the transpose of the connectivity matrix was used. Bidirectional transport was modeled by adding the spreading processes 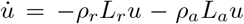, where *ρ*_*r*_ and *ρ*_*a*_ are the transport timescales of the retrograde and anterograde transport, and *L*_*a*_ = *L* and *L*_*r*_ = *L*^*T*^ are the graph Laplacians of anterograde and retrograde transport, respectively. Euclidean transport was modeled by substituting the anatomical connectivity matrix with a symmetric distance-based weight matrix as in previous work [17].

#### DIFF-R: Spreading and rise

Prion-like spreading is characterized not only by diffusive spreading but also by local aggregation processes. A commonly used model for prion-like aggregation is the heterodimer system,

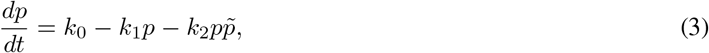

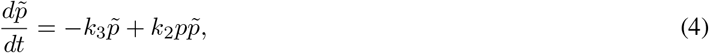

where *k*_0_ *>* 0 is the protein production rate, *k*_1_ *>* 0 and *k*_3_ *>* 0 are clearance rates, and *k*_2_ *>* 0 is the conversion rate from healthy to misfolded protein [19]. For 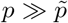, this system can be approximated by FKPP dynamics for the toxic protein 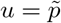,

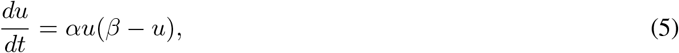

Where 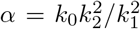denotes the global aggregation timescale and *β* = *α*^*−*1^(*k*_0_*k*_2_ *− k*_1_*k*_3_) is the carrying capacity.

We then obtain the spreading-and-rise model

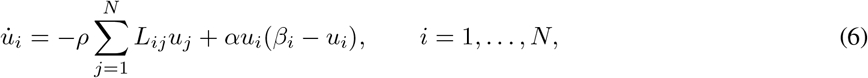

where *β*_*i*_ is the region-specific carrying capacity.

#### DIFF-RF: Spreading, rise, and fall

To capture the rise-and-fall pattern observed in the pathology data, we introduce a dynamic carrying capacity that decreases in response to local pathology burden:

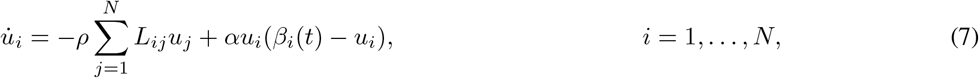

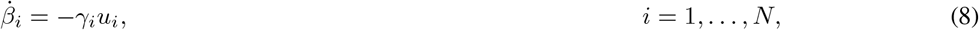

where *γ*_*i*_ *≥* 0 is the region-specific fall parameter. This additional process allows the local carrying capacity to decrease over time and produces rise-and-fall dynamics in pathology burden. A similar system was proposed to study dynamic clearance mechanisms in prion-like spreading models [69].

In the DIFF-R model, *β*_*i*_ is a region-specific parameter representing the local carrying capacity. In the DIFF-RF model, *β*_*i*_(*t*) is a state variable. Model inference for DIFF-RF is therefore performed on the initial condition *β*_*i*_(0), which we denote by *β*_*i*_ for notational simplicity.

### MCMC Bayesian estimation of modeling parameters

We inferred the parameters of the mathematical models using Bayesian MCMC, which has been used previously to parameterize prion-like propagation models [21, 22, 25, 70, 71]. The prior distributions are summarized in Table 2.

**Table 2:** Prior distributions for model parameters. The index *i* = 1, …, *N* represents brain regions. *N* (*µ, σ*) denotes a normal distribution and *N* ^+^(*µ, σ*) denotes a truncated normal distribution in (0, *∞*).

| Model | Parameter(s) | Prior |
| --- | --- | --- |
| DIFF | $\rho$ | $\mathcal{N}^+(50, 10)$ |
| DIFF-R | $\rho$ | $\mathcal{N}^+(0, 0.1)$ |
| | $\alpha$ | $\mathcal{N}^+(0, 0.1)$ |
| | $\beta_i$ ($i = 1, \dots, N$) | $\mathcal{N}(0, 1)$ |
| DIFF-RF | $\rho$ | $\mathcal{N}^+(0, 0.1)$ |
| | $\alpha$ | $\mathcal{N}^+(0, 0.1)$ |
| | $\beta_i$ ($i = 1, \dots, N$) | $\mathcal{N}(0, 1)$ |
| | $\gamma_i$ ($i = 1, \dots, N$) | $\mathcal{N}^+(0, 0.1)$ |

The initial condition is set with all regions set to zero, except for a set of initial seeding regions for which we infer initial values. For the striatal dataset, the initial seeding region was the ipsilateral striatum region, while for the hippocampal dataset the initial seeding sites were the ipsilateral CA1, CA3, and dentate gyrus.

As the DIFF model conserves total mass, we need a prior which is able to support a large enough initial condition to have enough mass to distribute throughout the network. This is not necessary in DIFF-R and DIFF-RF. As such, the prior for the initial conditions in the DIFF model is considerably wider than in the DIFF-R and -RF models. Additionally, the injection region(s) were omitted from the log-likelihood, as this is also needed to allow the DIFF model to find a posterior initial condition with a large enough mass to allow brain-wide propagation. For fair comparison across models, this exclusion was also applied to DIFF-R and DIFF-RF.

Let *y*_*ijt*_ denote the observed pathology burden for sample *j* in region *i* at timepoint *t*, and let *u*_*i*_(*t*, ***θ***) denote the corresponding model solution, where ***θ*** denote the model parameters and initial conditions. We assumed a normal likelihood with shared observation noise *σ*^2^,

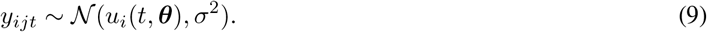

Posterior distributions were estimated using the No-U-Turn Sampler implemented in the Turing.jl package in the Julia programming language, with 1000 warmup iterations, 1000 posterior samples, target acceptance ratio 0.65 with four chains run in parallel [72]. Convergence was assessed using the Gelman–Rubin statistic 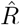 across chains (**Fig. S8**; hippocampal replication diagnostics in **Fig. S4**). For the models included in the main analyses, most parameters had 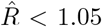, with a small number of exceptions. Prior–posterior comparisons for global parameters and pairwise comparisons among regional parameters are shown in **Fig. S9**. For the hippocampal DIFF-RF analysis, the lower temporal resolution of the dataset resulted in a less identifiable posterior than in the striatal analysis. We therefore used the striatal posterior distributions as priors for the global DIFF-RF parameters, while retaining the same priors for regional parameters as in the striatal model. Four chains were run with the same sampler settings as for the striatal analyses. Two converged to the same high-log-likelihood posterior mode, whereas the remaining chains occupied clearly lower-likelihood modes. The high-likelihood chains were therefore retained for the hippocampal vulnerability axis, transcriptomic, GSEA, and cell-type analyses. Chain-wise log-likelihoods and 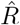 summaries are shown in **Fig. S4**.

### Model comparison

Mathematical models were compared using WAIC, AIC, BIC, mean squared error, and *R*^2^ relative to the observed mean [40, 73, 74]. WAIC was computed from pointwise log-likelihood values across posterior draws, treating each individual pathology measurement as a pointwise observation. WAIC was computed using 300 posterior draws per fitted model, as increasing the number of posterior draws beyond 300 did not substantially change WAIC estimates. Lower WAIC values indicate better expected predictive performance after accounting for effective model complexity. For reporting and visualization, WAIC values were expressed as ΔWAIC relative to the best-performing model within each comparison set. Uncertainty in ΔWAIC was estimated from paired pointwise WAIC differences across observations. For comparison, we also computed AIC and BIC, which compare model fit while penalizing parameter number, with BIC applying a stronger penalty for model complexity.

Model agreement was assessed using maximum *a posteriori* parameter estimates. The injection region data points were excluded, as these data points were not used to estimate model parameters. Mean squared error was computed between observed and model-predicted pathology values across region–timepoint observations. We also computed an identity-line *R*^2^, defined as 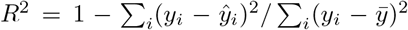, where *y*_*i*_ denotes observed pathology and ŷ_*i*_ denotes the corresponding model prediction. This metric quantifies improvement over the observed mean while evaluating predictions on their original scale, without fitting an additional regression slope or intercept.

### Connectivity and seeding null models

To assess whether model performance depended on the empirical connectome and injection location, we compared the fitted DIFF-RF model with two null configurations. For the connectivity null, the anatomical connectivity matrix was randomized by iterating over existing nonzero edges and assigning each edge weight to a randomly selected source– target pair. For the seeding null, the initial seeding region was randomized while retaining the empirical anatomical connectivity matrix. For each null configuration, the model was fit using the same inference procedure as for the empirical model and compared using WAIC.

### Gene–parameter association and enrichment analysis

To interpret the biological processes associated with the regional rise and fall parameters, we analyzed associations between regional gene expression and the inferred parameter maps from the DIFF-RF model. Analyses were performed using all regions with matched transcriptomic and model-parameter data. For each gene *g*, we fit a multiple linear regression model across regions, indexed by *i*, of the form

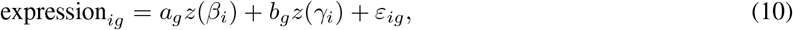

where *z*(*β*_*i*_) and *z*(*γ*_*i*_) denote standardized regional parameter values, (*a*_*g*_, *b*_*g*_) are gene-specific regression coefficients, and *ε*_*ig*_ is the residual error term. To make our results comparable across datasets, the parameters are *z*-transformed across the set of regional parameters from both datasets put together. This regression was performed across hemisphere-specific observations, yielding one coefficient pair per gene. Each gene is therefore represented as a point in the two-dimensional coefficient space (*a*_*g*_, *b*_*g*_).

To summarize the dominant structure of this coefficient space, we performed PCA across genes. Specifically, PCA was applied to the matrix of gene coefficients [(*a*_*g*_, *b*_*g*_)], treating genes as observations and regression coefficients as variables. The first principal component defined a scalar regional axis

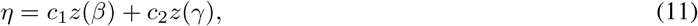

where (*c*_1_, *c*_2_) are the loadings of the first principal component. For each gene, we then computed the Pearson correlation between its regional expression profile and *η*. Genes were ranked by this correlation coefficient, which quantifies alignment with the vulnerability axis. Gene set enrichment analysis was performed using pre-ranked GSEA (`gseapy`) with the KEGG mouse pathway library (`KEGG_2019_Mouse`) [75]. Gene sets containing between 10 and 500 genes were included, and enrichment significance was assessed using 1000 permutations to obtain NES. Rather than focusing on enrichment significance alone, we examined the distribution of gene-level correlations within each pathway, allowing visualization of how pathway-associated genes align with the vulnerability axis.

For higher-level interpretation, KEGG pathways were grouped into broad functional categories reflecting biological processes commonly implicated in neurodegeneration (metabolism, protein homeostasis, synaptic function, neurodegenerative disease, and other). Assignment was performed using keyword-based matching of pathway names (Table S2). Over-representation of significantly enriched pathways within each category was assessed using Fisher’s exact test, with Benjamini-Hochberg correction applied to control for multiple comparisons.

To examine the directionality of the energy-metabolism signal, we curated genes encoding core TCA-cycle enzymes and oxidative phosphorylation complexes I–V (Table S1). For each dataset and module, we tested whether gene-level Pearson correlations with *η* were greater than zero using a one-sided one-sample *t*-test. The same curated genes were used to compare gene-level correlations across datasets using Spearman correlation.

### Cell-type composition analysis

To relate regional cell-type composition to the inferred model parameters, we analyzed associations between neurotransmitter class fractions and the regional dynamical parameters of the DIFF-RF model. Because cell-type fractions are compositional, fractions were transformed using the centered log-ratio (CLR) transform prior to statistical analysis [76]. Cell-type associations were evaluated primarily with respect to the regional axis *η* defined by the PCA described above, which summarizes the dominant joint variation of the rise and fall parameters. As for the gene-parameter association analysis, analyses were performed using all regions with matched cell-type and model-parameter data.

Spearman correlations were computed between CLR-transformed cell-type components and the regional values of *η*. Hemisphere-specific parameter estimates were retained as separate observations, while cell-type predictors were shared across hemispheres for each region.

Statistical significance was assessed using permutation tests that preserve the mapping between hemispheres belonging to the same region. In each permutation, region identities were shuffled while maintaining the shared predictor values for the two hemispheres of each region. Empirical *p*-values were obtained from 10,000 permutations. The false positive rate due to multiple testing across cell types was controlled using the Benjamini–Hochberg FDR.

To test whether monoaminergic populations were specifically associated with the vulnerability axis *η*, we defined a monoaminergic score for each region as the mean of the CLR-transformed fractions of dopaminergic, noradrenergic, serotonergic, and histaminergic neurons.

## Acknowledgements

We thank the Van Andel Institute Pathology and Biorepository Core (RRID:SCR 022912) for their assistance with tissue sectioning.

## Data Accessibility

Code used for this study is available at https://github.com/gretarsson/PrionNetworkModels/tree/main/paper-rf and archived on Zenodo at https://doi.org/10.5281/zenodo.20766155. Experimental data are archived on Zenodo at https://doi.org/10.5281/zenodo.21046224. Previously published striatal pathology and PANGEA transcriptomic data used in this study are available at https://doi.org/10.5281/zenodo.20648167 [17]. Posterior inference chains are archived on Zenodo at https://zenodo.org/records/21981450. Initial conversion of Nutil files to matrix output and grid heatmap generation: https://github.com/DaniellaDeWeerd/NutilToUsable (RRID:SCR 024753). Brain heatmaps and differential analysis of area occupied: https://github.com/vari-bbc/Mouse_Brain_Heatmap.

## Ethics

The authors declare that they have no competing interests.

