## Supplementary information for "Rise-and-fall dynamics reveal a molecular and cellular vulnerability axis in prion-like α-synuclein propagation"

### Supplementary information for Rise-and-fall dynamics reveal a molecular and cellular vulnerability axis in prion-like $\alpha$-synuclein propagation

Christoffer G. Alexandersen[1,2], Julia K. Brynildsen[1,2], Alice Prigent[2,3], Massimiliano Tamborrino[4], Anastasia Mantziou[4], Kevin Kurgat[2,3], Michael X. Henderson[2,3], and Dani S. Bassett[1,2,5,6]

[1]Department of Bioengineering, School of Engineering and Applied Science, University of Pennsylvania, PA, USA
[2]Aligning Science Across Parkinson's (ASAP) Collaborative Research Network, Chevy Chase, MD
[3]Department of Neurodegenerative Science, Van Andel Institute, Grand Rapids, MI, USA
[4]Department of Statistics, University of Warwick, Warwick, UK
[5]Department of Biomedical Engineering, Yale University, New Haven, CT, USA
[6]Department of Psychology, Yale University, New Haven, CT USA

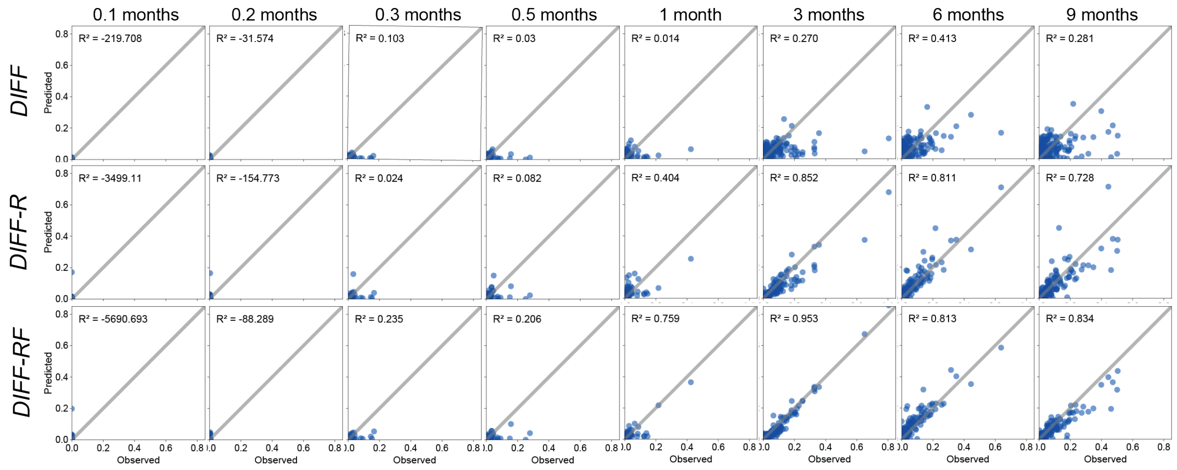


**Figure S1: Model predictive performance across time following striatal injection.** Predicted versus observed pathology at individual timepoints (all regions pooled). Panels correspond to increasing time after injection (0.1 to 9 months). The diagonal indicates identity.

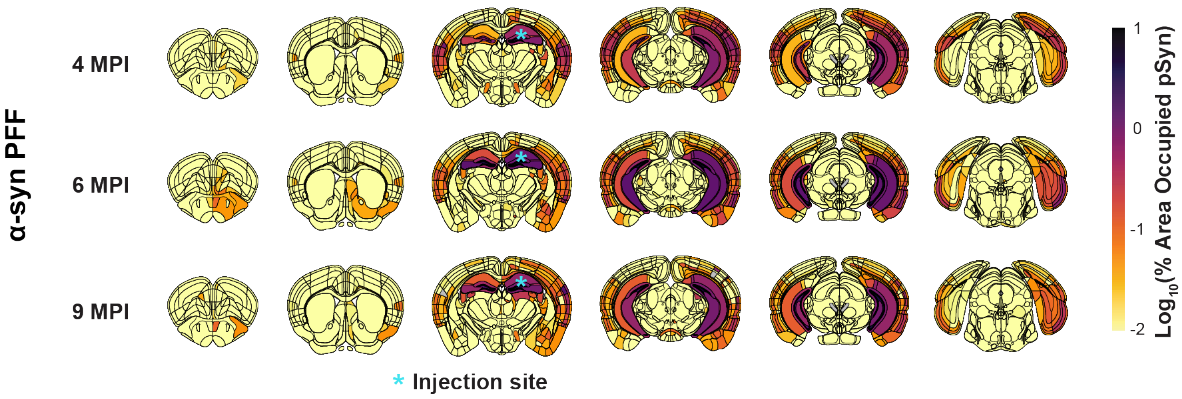


**Figure S2: $\alpha$-Synuclein pathology distribution following hippocampal injection of $\alpha$-synuclein PFFs.** Wildtype mice injected with $\alpha$-synuclein PFFs in the hippocampus and overlaying cortex were aged 4, 6, or 9 MPI. Anatomical heatmaps showing the log-transformed percentage of area occupied by phosphorylated $\alpha$-synuclein. The injection site is indicated by an asterisk.

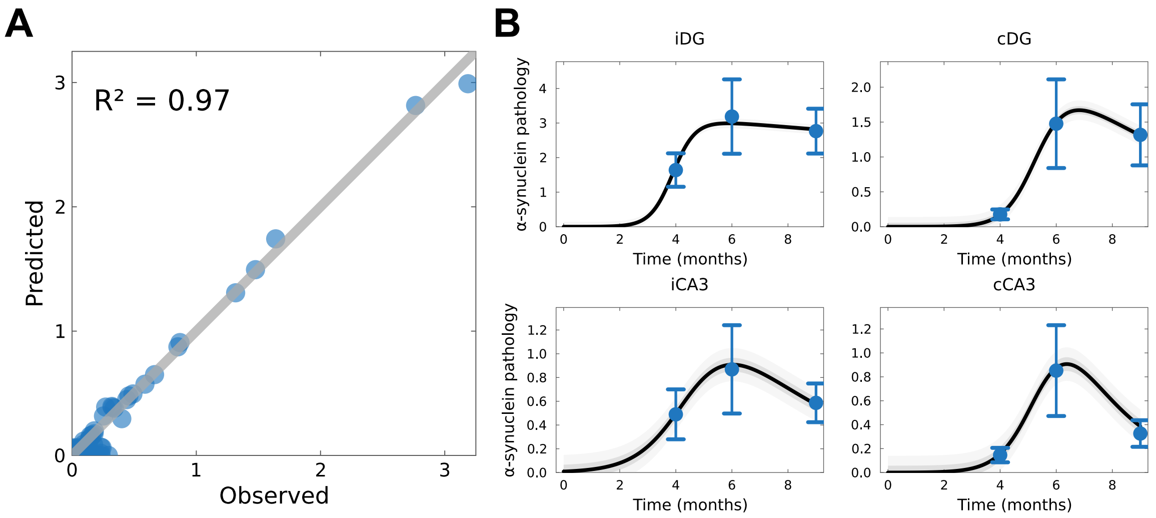


**Figure S3: Model performance of DIFF-RF in the hippocampal dataset.** (**A**) Predicted versus observed $\alpha$-synuclein pathology for the hippocampal dataset, pooling all regions and timepoints. Each point corresponds to a region–timepoint pair; the diagonal indicates identity. (**B**) Posterior mean retrodictions for the four regions with highest observed pathology. Blue points and error bars show observed mean $\pm$ SE, black curves show model predictions, and gray bands show posterior predictive uncertainty.

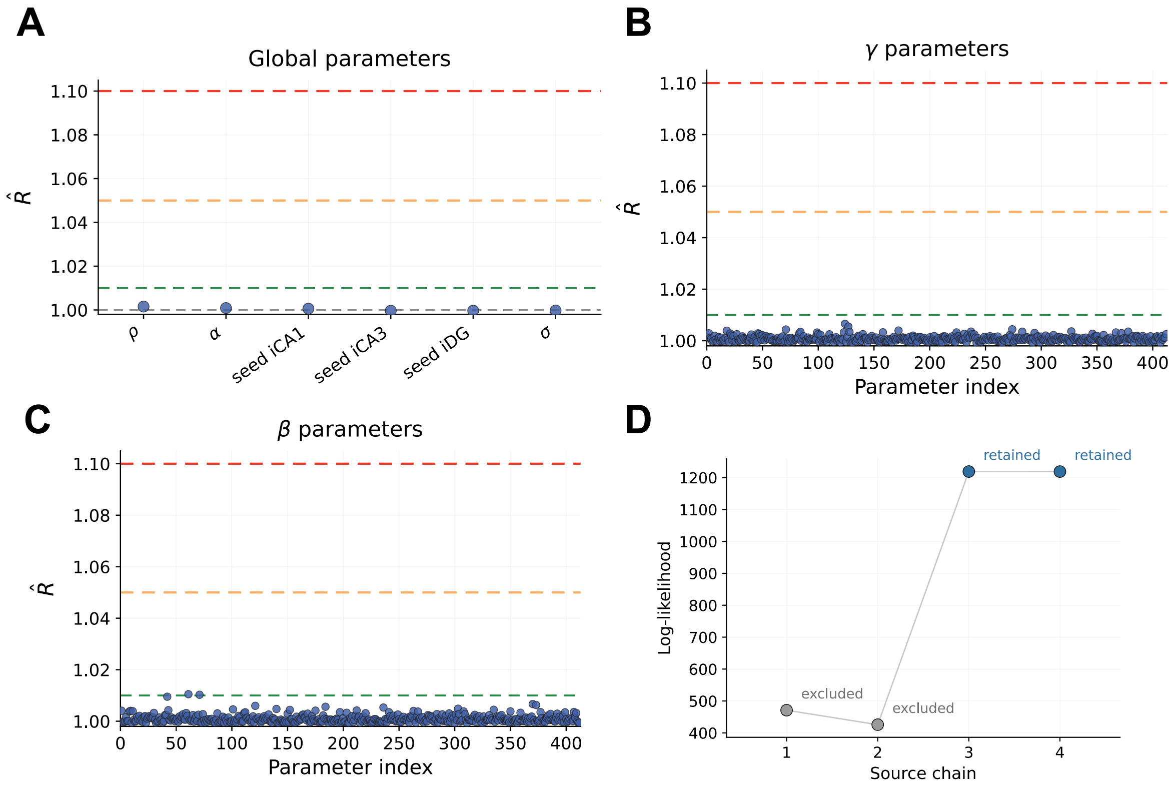


**Figure S4: Posterior diagnostics for the hippocampal DIFF-RF model.** (**A–C**) Gelman–Rubin statistics ($\hat{R}$) for global, fall ($\gamma$), and rise ($\beta$) parameters. Horizontal dashed lines indicate common convergence thresholds ($\hat{R} = 1.01$, 1.05, and 1.10). (**D**) Chain-wise log-likelihood evaluated over all observations. The retained chains show high and concordant log-likelihood, whereas excluded chains occupy lower-likelihood modes.

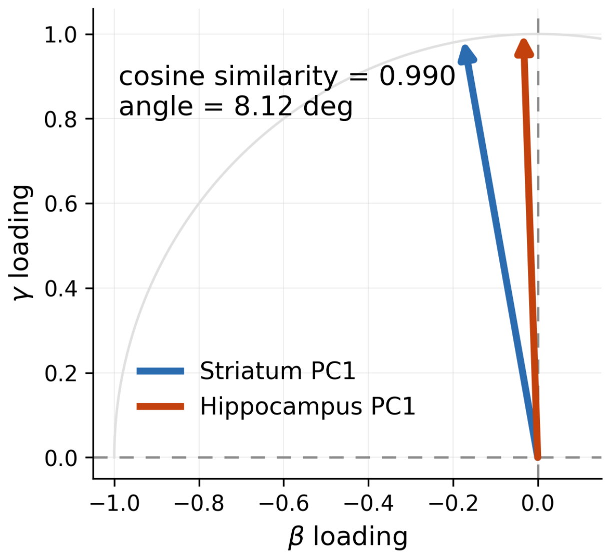


**Figure S5: Vulnerability axes comparison across datasets.** Comparison of the PC1 / vulnerability axes found in the striatal and hippocampal datasets in raw $(\beta_i, \gamma_i)$ coefficient space.

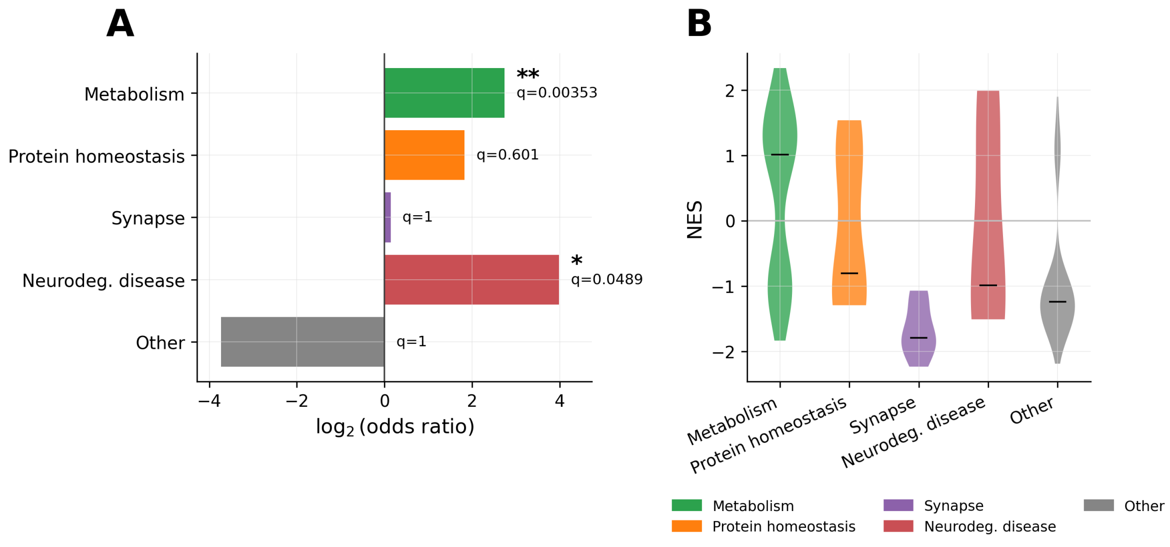


**Figure S6: Additional analyses of gene enrichment for the hippocampal dataset.** (**A**) Enrichment of functional gene categories along the vulnerability axis, shown as $\log_2$ odds ratios with Benjamini–Hochberg adjusted $q$-values. (**B**) Distribution of normalized enrichment scores (NES) across functional categories.

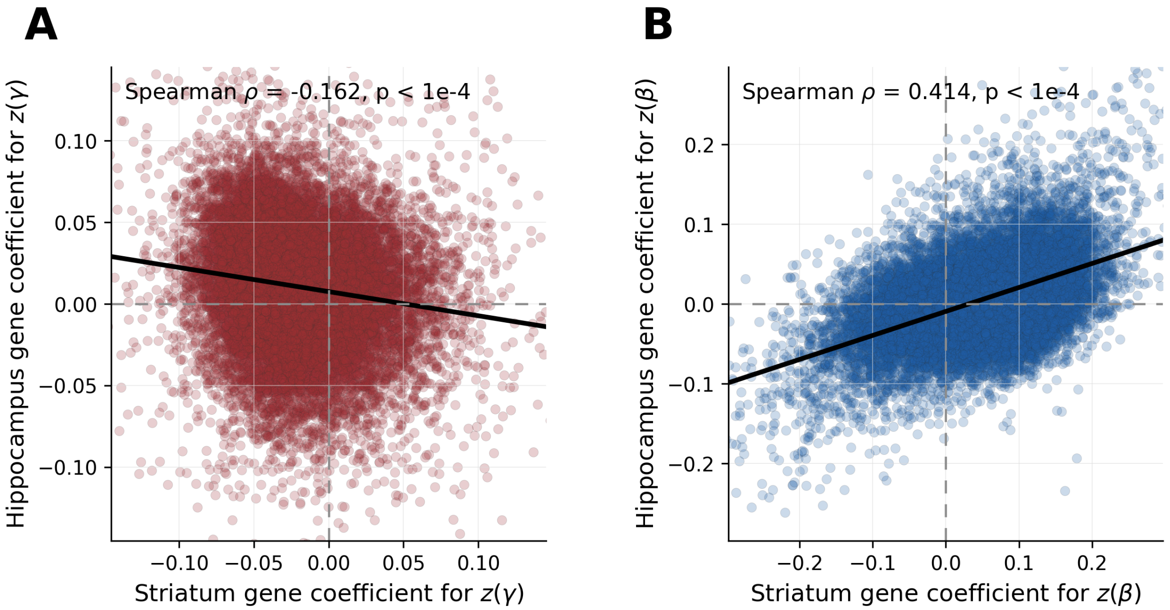


**Figure S7: Reproducibility of gene-parameter associations across datasets.** (**A**) Comparison of gene-level multiple-regression coefficients in the striatal and hippocampal datasets for the fall parameter $z(\gamma)$. (**B**) Same as in panel (A) but for the rise parameter $z(\beta)$.

**Table S1: Curated energy-metabolism genes used to assess directionality of the metabolic vulnerability-axis signature.** Genes were selected as core TCA-cycle enzymes or nuclear-encoded oxidative phosphorylation complex subunits and were analyzed as module-level groups.

| Module | Genes | Role |
|---|---|---|
| TCA cycle | Aco2, Cs, Dld, Dlst, Fh1, Idh3a, Idh3b, Idh3g, Mdh2, Ogdh, Ogdhl, Sdha, Sdhb, Sdhc, Sdhd, Sucla2, Suclg1, Suclg2 | Core TCA-cycle enzymes carrying carbon flux and generating reducing equivalents. |
| Complex I | Ndufa1, Ndufa10, Ndufa11, Ndufa12, Ndufa13, Ndufa2, Ndufa3, Ndufa4, Ndufa5, Ndufa6, Ndufa7, Ndufa8, Ndufa9, Ndufab1, Ndufb1, Ndufb10, Ndufb11, Ndufb2, Ndufb3, Ndufb4, Ndufb5, Ndufb6, Ndufb7, Ndufb8, Ndufb9, Ndufc1, Ndufc2, Ndufs1, Ndufs2, Ndufs3, Ndufs4, Ndufs5, Ndufs6, Ndufs7, Ndufs8, Ndufv1, Ndufv2, Ndufv3 | NADH:ubiquinone oxidoreductase subunits; NADH oxidation and proton pumping. |
| Complex II | Sdha, Sdhb, Sdhc, Sdhd | Succinate dehydrogenase; oxidizes succinate and feeds electrons into ubiquinone. |
| Complex III | Cyc1, Uqcr10, Uqcr11, Uqcrb, Uqcrc1, Uqcrc2, Uqcrfs1, Uqcrh, Uqcrq | Cytochrome bc1 complex; transfers electrons from ubiquinol toward cytochrome c. |
| Complex IV | Cox4i1, Cox4i2, Cox5a, Cox5b, Cox6a1, Cox6a2, Cox6b1, Cox6b2, Cox6c, Cox7a1, Cox7a2, Cox7a2l, Cox7b, Cox7b2, Cox7c, Cox8a, Cox8b, Cox8c | Cytochrome c oxidase subunits; terminal electron transfer to oxygen and proton pumping. |
| Complex V | Atp5a1, Atp5b, Atp5c1, Atp5d, Atp5e, Atp5g1, Atp5g2, Atp5g3, Atp5h, Atp5j, Atp5j2, Atp5k, Atp5l, Atp5md, Atp5mpl, Atp5o, Atp5pb | ATP synthase subunits; converts proton-motive force into ATP. |

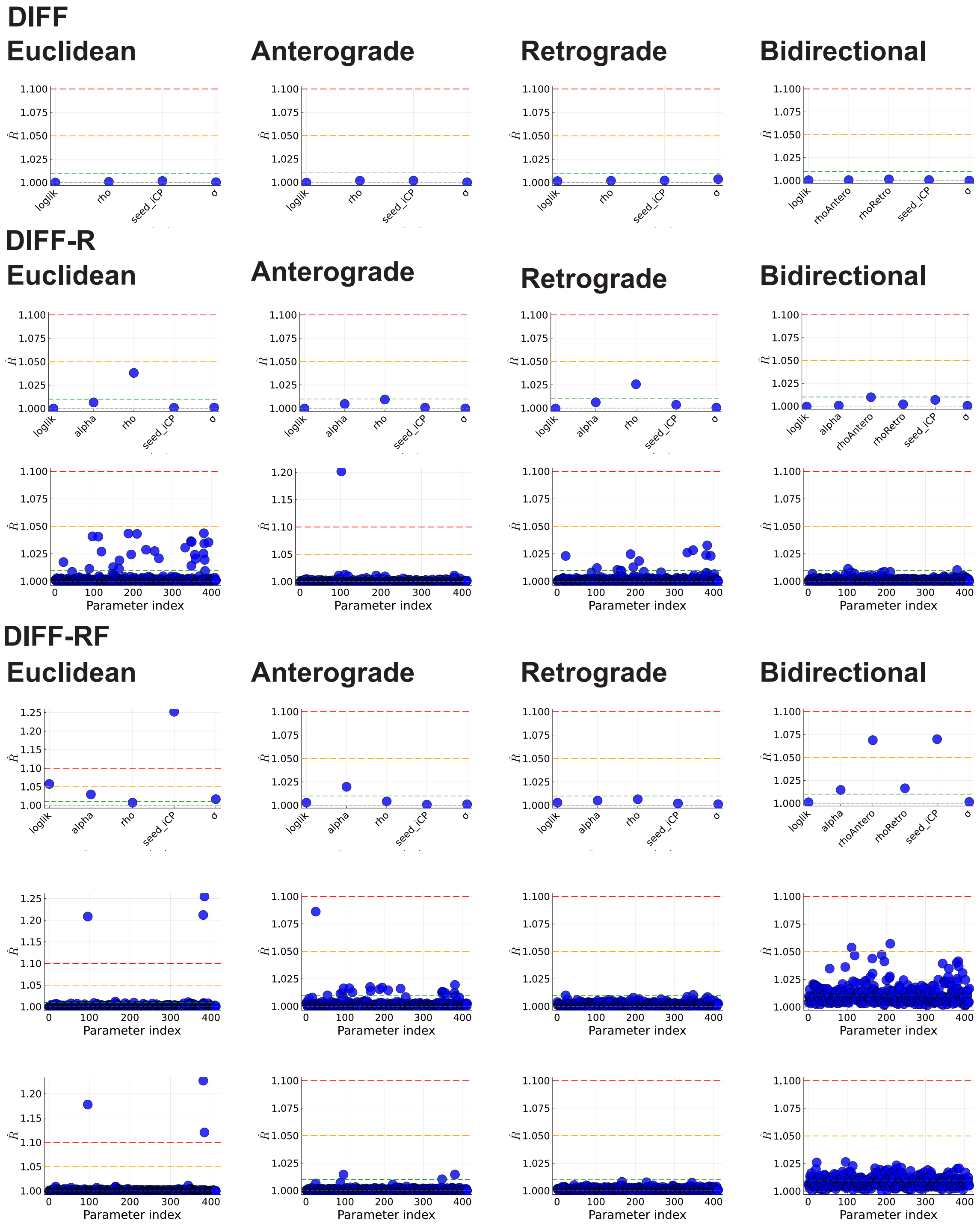


**Figure S8: Convergence diagnostics across models for the striatal dataset.** Gelman–Rubin statistics ($\hat{R}$) for all inferred parameters across model variants and datasets. Each panel shows $\hat{R}$ values either aggregated by parameter type (top rows) or for all individual parameters indexed sequentially (rows labeled "Parameter index"). Horizontal dashed lines indicate common convergence thresholds ($\hat{R} = 1.01$, 1.05, and 1.10).

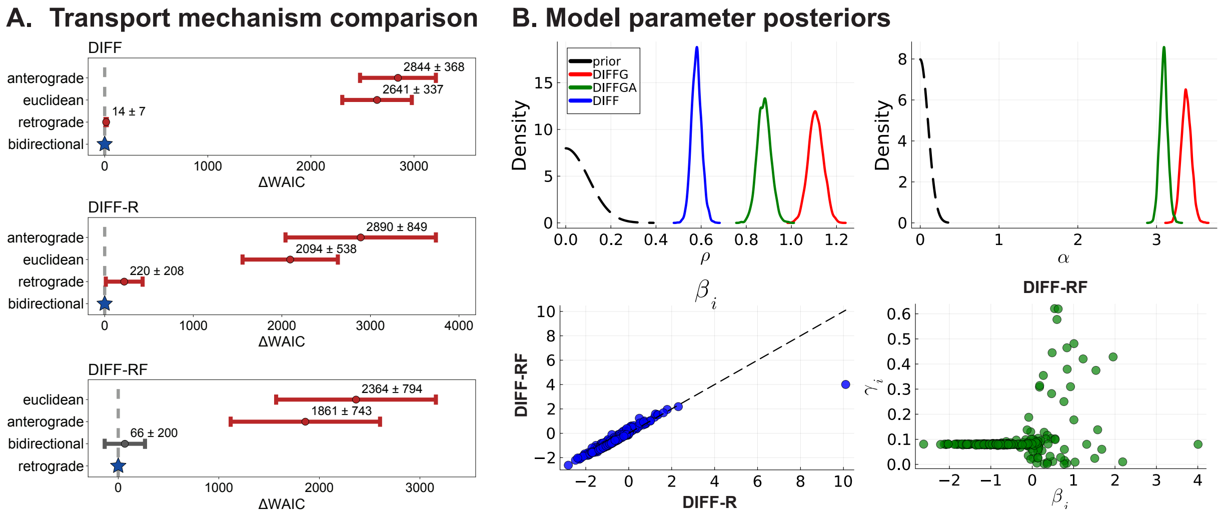


**Figure S9: Prior and posterior distributions and regional parameter relationships for the striatal dataset.** (A) Prior (black dashed) and posterior densities for the global transport parameter $\rho$ across model classes (DIFF, DIFF-R, DIFF-RF). (B) Prior (black dashed) and posterior densities for the global growth parameter $\alpha$ in DIFF-R and DIFF-RF. (C) Comparison of the posterior means of the regional carrying capacity parameters $\beta_i$ inferred under DIFF-R and DIFF-RF; the dashed line denotes identity. (D) Means of the posterior regional decay parameters $\gamma_i$ versus carrying capacity parameters $\beta_i$ in DIFFGA.

**Table S2: Keyword-based mapping used to assign KEGG pathways to broad functional categories for summary analyses.** Pathways were matched case-insensitively to the first category whose keyword list was satisfied; pathways that matched none of the listed keywords were assigned to *Other*.

| **Category** | **Keywords used for assignment** |
|---|---|
| Metabolism | metabolism, metabolic, oxidative phosphorylation, mitochond, glycolysis, tca, citrate cycle, respiratory chain, fatty acid, lipid, cholesterol, biosynthesis, amino acid, nucleotide, energy |
| Protein homeostasis | proteasome, ubiquitin, ubiquitination, autophagy, lysosome, endoplasmic reticulum, er , chaperone, protein folding, protein processing, ribosome, translation, mrna, degradation, unfolded protein, quality control, mitophagy |
| Synaptic function | synapse, synaptic, dopaminergic synapse, glutamatergic synapse, gabaergic synapse, neurotransmitter, vesicle, long-term potentiation, long-term depression |
| Neurodegenerative disease | parkinson, alzheimer, huntington, prion, amyotrophic, neurodegenerative, als |
| Other | Any pathway name not matching the keyword lists above |
